# Microsatellite break-induced replication generates highly mutagenized extrachromosomal circular DNAs

**DOI:** 10.1101/2024.01.12.575055

**Authors:** Rujuta Yashodhan Gadgil, S. Dean Rider, Resha Shrestha, Venicia Alhawach, David C. Hitch, Michael Leffak

## Abstract

Extrachromosomal circular DNAs (eccDNAs) are produced from all regions of the eucaryotic genome. In tumors, highly transcribed eccDNAs have been implicated in oncogenesis, neoantigen production and resistance to chemotherapy. Here we show that unstable microsatellites capable of forming hairpin, triplex, quadruplex and AT-rich structures generate eccDNAs when integrated at a common ectopic site in human cells. These non-B DNA prone microsatellites form eccDNAs by replication-dependent mechanisms. The microsatellite-based eccDNAs are highly mutagenized and display template switches to sister chromatids and to nonallelic chromosomal sites. High frequency mutagenesis occurs within the eccDNA microsatellites and extends bidirectionally for several kilobases into flanking DNA and nonallelic DNA. Mutations include mismatches, short duplications, longer nontemplated insertions and large deletions. Template switching leads to recurrent deletions and recombination domains within the eccDNAs. Template switching events are microhomology-mediated, but do not occur at all potential sites of complementarity. Each microsatellite exhibits a distinct pattern of recombination, microhomology choice and base substitution signature. Depletion of Rad51, the COPS2 signalosome subunit or POLη alter the eccDNA mutagenic profiles. We propose an asynchronous capture model based on break-induced replication from microsatellite-induced DNA breaks for the generation and circularization of mutagenized eccDNAs and genomic homologous recombination deficiency (HRD) scars.

## Introduction

DNA replication is continuously challenged by exogenous and endogenous replication stressors including DNA damage, tightly bound proteins, transcription complexes and non-B (noncanonical, non-Watson-Crick) DNA structures (1,2). The formation of non-B structures is a common feature of short (∼1-9 bp) tandemly repeated sequences termed microsatellites, which are highly abundant in metazoan genomes, representing approximately 3% of human DNA. The structural instability of microsatellites has been attributed to their tendency to form noncanonical structures, e.g. DNA hairpins, four-stranded G-quadruplex (G4) DNA, triple stranded Hoogstein DNA (H-DNA), and unwound or collapsed AT-rich DNA structures (3–8). Microsatellite instability has been causally linked to more than forty neurological and developmental disorders including Huntington’s disease (HD), Myotonic dystrophy type 1 (DM1), and Spinocerebellar ataxia (SCA) type 10 (3,9–13). Short tandem repeats are also found at chromosome fragile sites where they are hotspots of DNA double strand breaks (DSBs), mutagenesis and nonreciprocal template switches. Accordingly, triplex-and quadruplex-prone sequences co-localize with breakpoints in the human c-myc gene in Burkitt’s lymphoma and diffuse large B-cell lymphoma, and in the BCL-2 gene in follicular lymphoma (10,14–19).

Hairpin, AT-rich, triplex and G4 DNA are susceptible to replication-dependent DSBs, hypermutagenesis and chromosomal template switches (20–30) by the homology-dependent mechanisms of break-induced replication (BIR) and BIR-like long-tract gene conversion (20–22). Repetitive elements and regions of chromosome instability are also enriched in eccDNAs (14,31,32). In addition to homologous recombination (HR), nonhomologous end-joining (NHEJ) and mismatch repair (MMR) have been implicated in the production of eccDNA during breakage-fusion-bridge (BFB) cycles, circularization following chromothripsis, episome extrusion due to stalled replication, and through cycles of nonallelic template switches and excision (14,33–39), however, there is little mechanistic information regarding how specific microsatellites become configured into eccDNAs.

eccDNAs are double stranded, circular molecules, frequently with the ability to replicate autonomously (28,33,40–46). eccDNA is found in all eukaryotes, including human normal cells and tumors (31,40,45–49). Depending on their size or content eccDNAs can be classified as episomes or double minutes (DMs) (∼100 kb - 3 Mb), small polydisperse circular DNA (spcDNA) (∼100 bp -10 kb), and microDNA (∼100 - 400 bp) (14,40,50–52). eccDNAs have been associated with the etiology of cancer, neurological disorders, autoimmunity, aberrant cell signaling and ageing due to gene rearrangement/amplification, altered transcriptional activity and enhanced chromatin accessibility (34,42,52–56). Extrachromosomal DNAs in microglia have recently been reported to lead to neurodegeneration (57), raising the possibility that replication-dependent microsatellite instability in microglia may contribute to the degeneration of post-mitotic neurons in HD, DM1 and SCA10.

We have analyzed the formation of eccDNA from four different microsatellites (G4, H-DNA, hairpin, and AT-rich DNA) at the single molecule level, integrated at the same ectopic chromosomal site. Using inverse PCR (iPCR) we find that each of these microsatellites produces eccDNAs containing unique template switching events which are recurrent, nonrandom, and distinct from those of the other microsatellites. The structures of the eccDNAs are dependent on DNA replication, and the eccDNA recombinants are mutagenized at ∼1000-fold the wild type rate. The microsatellite repeats themselves are hotspots for mutagenesis, including deletions, insertions, and base substitutions. Repeat-induced mutagenesis extends more than 5-10 kb bidirectionally from each microsatellite. Template switching events abundant in eccDNAs are strongly directed by distinct patterns of microhomology, and occur within the ectopic site and to nonallelic chromosomes.

## Materials and Methods

### Cell culture

HeLa/406 acceptor cells containing a single FRT site were used for the construction of all cell lines used in this study (20–22,58,59). HeLa/406 cells were co-transfected using dual fluorescence (dTomato, eGFP) donor plasmids and the pOG44 expression vector that produces FLP recombinase (21,22). Clonal cell lines were derived by limiting dilution. Cells were maintained on DMEM supplemented with 10% calf serum, 1% penicillin-streptomycin and 5% CO_2_ at 37 °C. For replication stress experiments, cells were treated with hydroxyurea (HU, 0.2 mM, 4 days) or aphidicolin (APH, 0.2 uM, 2 days), after which the cells were returned to standard medium and cultured for an additional 4 days to allow the turnover of preexisting dTomato and eGFP proteins. In protein knockdown experiments, cells were transfected with siRNA (final concentration 50 nM) or shRNA (15 nM final concentration) once or twice over a 48 hr period, washed, and allowed to recover in fresh medium for 4 days.

Cell lines are named DF/myc for the dual fluorescence genes and the presence of the c-myc core origin, followed by the microsatellite of choice; DF/myc(G4), DF/myc(H3), DF/myc(CAG)_102_ and DF/myc(ATTCT)_47_. For simplicity the cell lines are referred to as G4, H3, (CAG)_102_, and (ATTCT)_47_, respectively. The sequences of the G4 and H3 microsatellites have been published (21). Flow cytometry was performed on a BD Accuri 6 flow cytometer. All flow experiments were calibrated against the same set of yellow (dTom^+^, eGFP^+^), green (dTom^-^, eGFP^+^), red (dTom^+^, eGFP^-^), and double negative (dTom^-^, eGFP^-^) stable marker cell lines. The ectopic microsatellite cells had undergone ∼150 (G4, H3) or ∼200 ((CAG)_102_, (ATTCT)_47_) generations from the time of transfection through limiting dilution, clonal outgrowth and flow cytometry/DNA isolation. The siRNA and shRNA sequences used for protein knockdown are shown in Supplementary Table 1.

### Inverse PCR

DNA extraction was performed using Tissue DNA kits (Omega, D3396-00S). Inverse PCR (iPCR) was performed on undigested total genomic DNA using Q5 HotStart polymerase (New England Biolabs, M0494S). The primers used for amplification are shown in Supplementary Table 2. PCR Products were electrophoresed on 0.8% agarose gels. The amplified products were purified (Omega, D6492-01) before PacBio Sequel IIe sequencing (Azenta, Plainfield, N.J).

### DNA sequence analysis

To decatenate the iPCR products in the circular consensus reads, bespoke shell scripts were used to identify and mark the termini of individual PCR products within the concatemer strings, based on the primer sequences used for PCR. Individual PCR product sequence strings were then extracted for mapping. Hybrid reference genomes for each ectopic site or iPCR sequence were generated which included HeLa-based sequences for each chromosome in the human reference genome (GRCh38) (27). BWA was used to index the reference genomes and BWA-MEM (60) was used for mapping sequence reads against the hybrid reference sequences containing the engineered ectopic site, or the expected iPCR product. Sequence reads that did not contain ectopic site sequences for at least 6 bp 3’ to each iPCR primer were considered non-specific PCR products and were removed from the next round of mapping.

Duplicated sequences were also identified and removed. The second round of mapping was performed on unique reads which contained ectopic site sequences. Samtools (61) was used to sort and generate binary output for the mapped datasets (.bam and .bai files). Samtools was also used to generate statistics on the mapped datasets such as depth of coverage, contig length vs. number of reads, deletion events, deleted nucleotides, insertion events, inserted nucleotides, base mismatches, and pileup of the data. Pileup data was parsed using a bespoke shell script to summarize the total number of insertions, deletions and mismatches at a given position.

Additionally, the perl script pileup2baseindel.pl (https://github.com/riverlee/pileup2base) was used to generate detailed information on indel sequences. The sam2paf.js script from minimap2 (https://github.com/lh3/minimap2) (62) was used to convert .sam files to .paf format for use as input for analysis by ALVIS (63). A bespoke script was used to extract the mapped locations of fragments from chimeric reads and data were formatted for use with Circos (64). Integrative Genomics Viewer (65) was used to visualize and understand the composition of reads, links between mapped regions, and alignments against the genome. BLAST was used for base composition analysis. The .sam files were sorted and converted to .bam format which was also indexed for use with Ribbon (66). The number of reads in each Ribbon alignment is shown in the corresponding figure legend. Student’s t-test for random microhomology length was performed against a random number array generated by Graphpad Prism (https://www.graphpad.com). Sequence alignments were also performed using Snapgene (https://www.snapgene.com) or NCBI BLAST (https://blast.ncbi.nlm.nih.gov).

## Results

### Model systems of microsatellite instability

Microsatellites expanded beyond approximately 40-50 tandem repeats are unstable at their natural loci and at ectopic sites in mammalian genomes (22,26,67–73). We hypothesized that instability at microsatellites capable of forming quadruplex, triplex, hairpin or AT-structured DNAs could lead to the formation of eccDNAs. To test this hypothesis, we derived cell lines containing microsatellite DNAs integrated at an ectopic site (ES) in HeLa cells, where these microsatellites had been shown to display replication-dependent instability (20,21,26,59,67,74–76). The non-B DNA microsatellite cell lines were each generated by single copy integration at the same ectopic FLP recombinase site, alongside a copy of the 2.4 kb c-myc core replication origin (21,22,58,59) (Figure 1). The ES c-myc origin is active in >90% of replications, and displays the chromatin structure, replication protein binding, and early S-phase replication timing pattern of the endogenous c-myc origin (22,30,59,67,77). Dual fluorescence (DF; dTomato (dTom) and eGFP) reporter genes were positioned within the ES to allow DSBs to be detected by flow cytometry based on the loss of reporter protein fluorescence. AluYa5/IVS sequences derived from the UBE2T locus provided targets for homology-dependent recombination (78).

**Figure 1.**
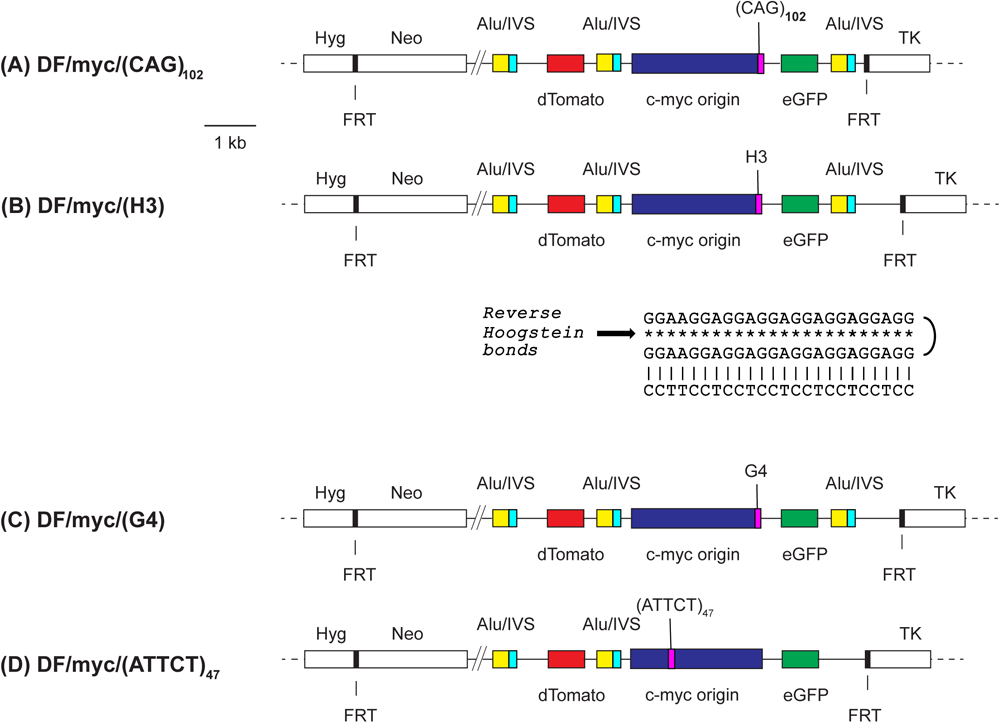
Ectopic site maps. Microsatellite sequences (A) (CAG)102, (B) above; H3 (21) (triplex forming); below, predicted triplex conformation; (C) G4 (quadruplex forming) (21), and; (D) (ATTCT)47 (30) were integrated at the FLP recombinase target (FRT) site in HeLa/406 cells (177) (CAG)102, H3 and G4 were placed in the lagging template strand; (ATTCT)47 was placed in the upper (5’-3’) strand template of the 2.4 kb c-myc core origin (30,177). Alu/IVS sequences were derived from the UBE2T locus (78). Hyg, hygromycin resistance; Neo, neomycin resistance; TK, HSV thymidine kinase; FRT, FLP recombinase target.

Each cell line is named for the non-B DNA in the lagging strand template when replicated from the c-myc origin. Triplex (H-DNA, H3) and quadruplex (G4) inserts were derived from the PKD1 IVS21 homopurine-homopyrimidine asymmetric mirror repeat (79). The H3 microsatellite (21) was designed to fulfill the requirements of homopurine-homopyrimidine mirror-symmetry (80,81) (Figure 1B).

The canonical G4 consensus sequence is (G)_3_(N)_1-7_(G)_3_(N)_1-7_(G)_3_(N)_1-7_(G)_3_, although consensus matches with loop regions as long as (N)_30_ have been reported to form stable G4 structures (82–84). The ectopic G4 microsatellite contained five matches to the canonical consensus in the lagging strand template of the c-myc origin (21).

The (CAG)_102_ insert length was chosen to exceed the threshold associated with the DM1 phenotype (85), and has been shown to adopt a hairpin structure in vitro and in vivo (59,67). The (ATTCT)_47_ microsatellite has been shown to restore replication initiation activity to the core c-myc origin inactivated by deletion of the DNA unwinding element (DUE), and to induce replication-dependent instability (30).

The non-B DNA cell lines are referred to hereafter by their ES non-B DNA ((CAG)_102_, G4, H3, and (ATTCT)_47_). Each clonal cell line was derived by limiting dilution. Upon clonal outgrowth, mixed flow cytometry profiles (dTom^+^, eGFP^+^; dTom^+^, eGFP^-^; dTom^-^, eGFP^+^; dTom^-^, eGFP^-^) were displayed by (CAG)_102_, G4, H3 and (ATTCT)_47_ cells (Figure 2), indicating that the same, or different, microsatellite sequences can undergo dissimilar forms of mutagenesis (86,87).

**Figure 2.**
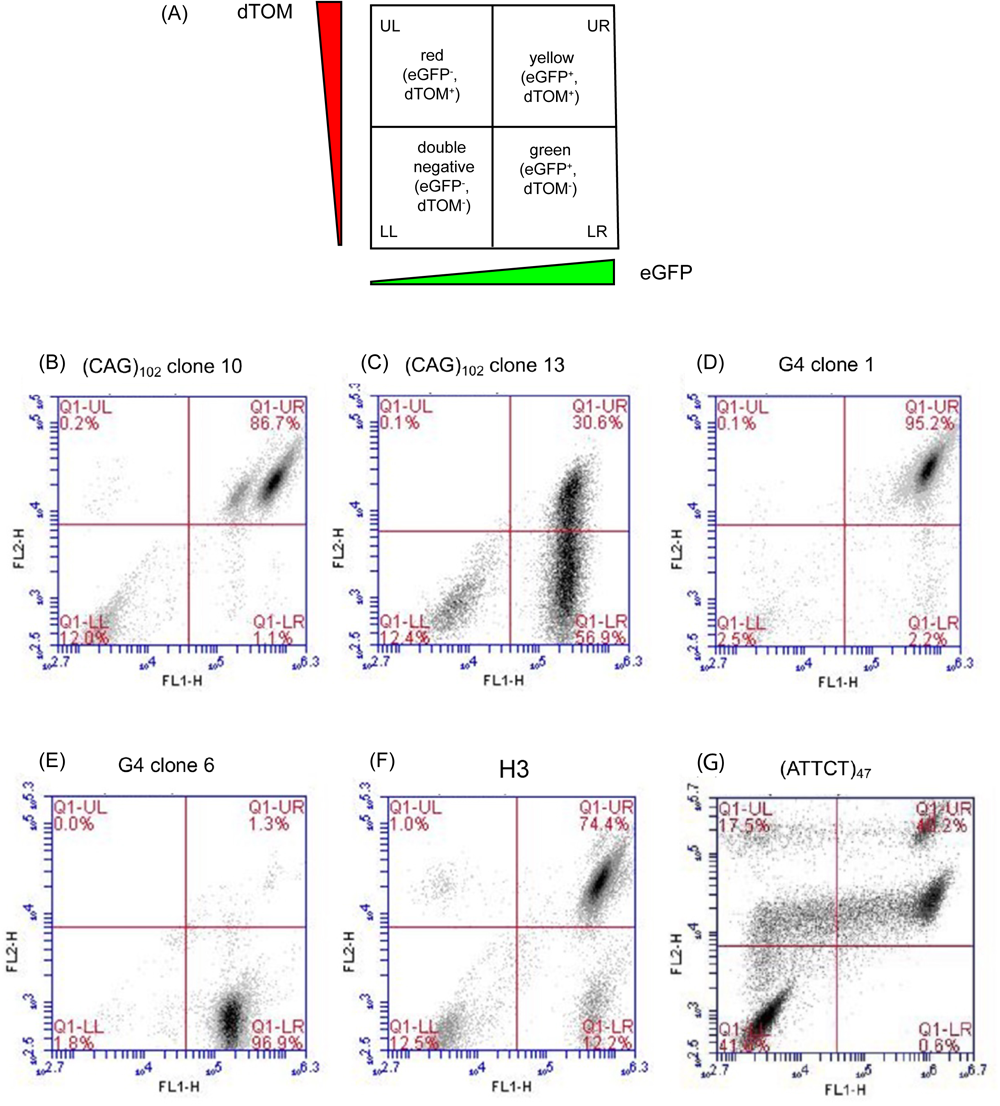
Flow cytometry analysis of microsatellite instability. (A) Schematic key indicating red (dTomato+, eGFP-), yellow (dTomato+, eGFP+), green (dTomato-, eGFP+), and double negative (dTomato-, eGFP-) cells; (B) G4 clone 1; (C) G4 clone 6; (D) (ATTCT)47; (E) H3; (F) (CAG)102 clone 13; (G) (CAG)102 clone 10.

(CAG)_102_ clone 10 (c.10) cells contained two yellow cell populations (dTom^+^, eGFP^+^; upper right (UR) quadrant) differing in the intensity of eGFP fluorescence, whereas more than 50% of the (CAG)_102_ clone 13 (c.13) cells lost expression of the dTom marker upstream of the (CAG)_102_ microsatellite. DNA sequencing across the (CAG)_102_ c.13 repeat (see below) showed the full length (CAG)_102_ microsatellite and contracted repeats with fewer than 30 (CAG) triplets.

In the two G4 clones, G4 clone #1 (G4 c.1), had lost half of the G4 insert and G4 clone #6 (G4 c.6) had lost the entire quadruplex consensus sequence (21). Despite having undergone a partial microsatellite deletion, G4 c.1 retained both fluorescent reporter genes and appeared in the upper right (Figure 2(D); UR, yellow) flow cytometry quadrant. In contrast, most G4 c.6 cells (Figure 2(E)) had lost or mutated the dTom marker and appeared in the LR green quadrant.

The H3 cells retained the triplex DNA insert during clonal outgrowth (21) but showed modest instability after long term culture, indicated by the appearance of green cells (dTom^-^, eGFP^+^; LR quadrant), and double negative (dTom^-^, eGFP^-^) cells (Figure 2(F)). In contrast to the G4 or (CAG)_102_ cells, the H3 cells also generated a small percentage (∼1%) of red cells.

The flow cytometry pattern of the (ATTCT)_47_ cells (Figure 2(G)) was substantially different from those of the other five cell clones. Two populations of yellow cells were evident, differing in the intensity of the dTom signal. We speculate that, as in (CAG)_102_ c.10 cells, the more highly fluorescent (ATTCT)_47_ cells contain extrachromosomal copies of the ES (but with mutations in the dTom reporter). Similarly, we propose that the less highly fluorescent (CAG)_102_ c.10 population contains extrachromosomal copies of the ES with mutations in the eGFP reporter. Both populations of yellow (ATTCT)_47_ cells also appeared to generate red cell populations (UL quadrant) consistent with loss of eGFP expression by local mutation or recombination, and double negative cells (LL quadrant) indicating loss of expression of both reporter genes in the same cell.

All of the flow cytometry profiles shown were derived from cells under unperturbed growth conditions, indicating that microsatellite instability occurs in the presence of endogenous replication stress.

### Microsatellites generate eccDNAs

To test the hypothesis that microsatellite DNAs could generate eccDNAs we used inverse PCR (iPCR) on undigested total genomic DNA (20,88–90). iPCR primers were designed that face away from each other at the ES, to amplify only circular templates. The iPCR products ranged from ∼250 bp to ∼10 kb (Figure 3). The maximum size of the observed iPCR products may reflect a limitation on the extent of the iPCR by Q5 DNA polymerase. We note as well that these products do not reveal the length of the DNA between the 5’ ends of the iPCR primers. Each of the microsatellite-containing ES’s generated distinct iPCR products; in contrast the empty ES, which contains the active c-myc origin core but no additional non-B DNA, did not produce clear iPCR products (Figure 3, lane 14). We conclude that each microsatellite represents a hotspot for eccDNA formation, and generates multiple preferred, distinct recombinants.

**Figure 3.**
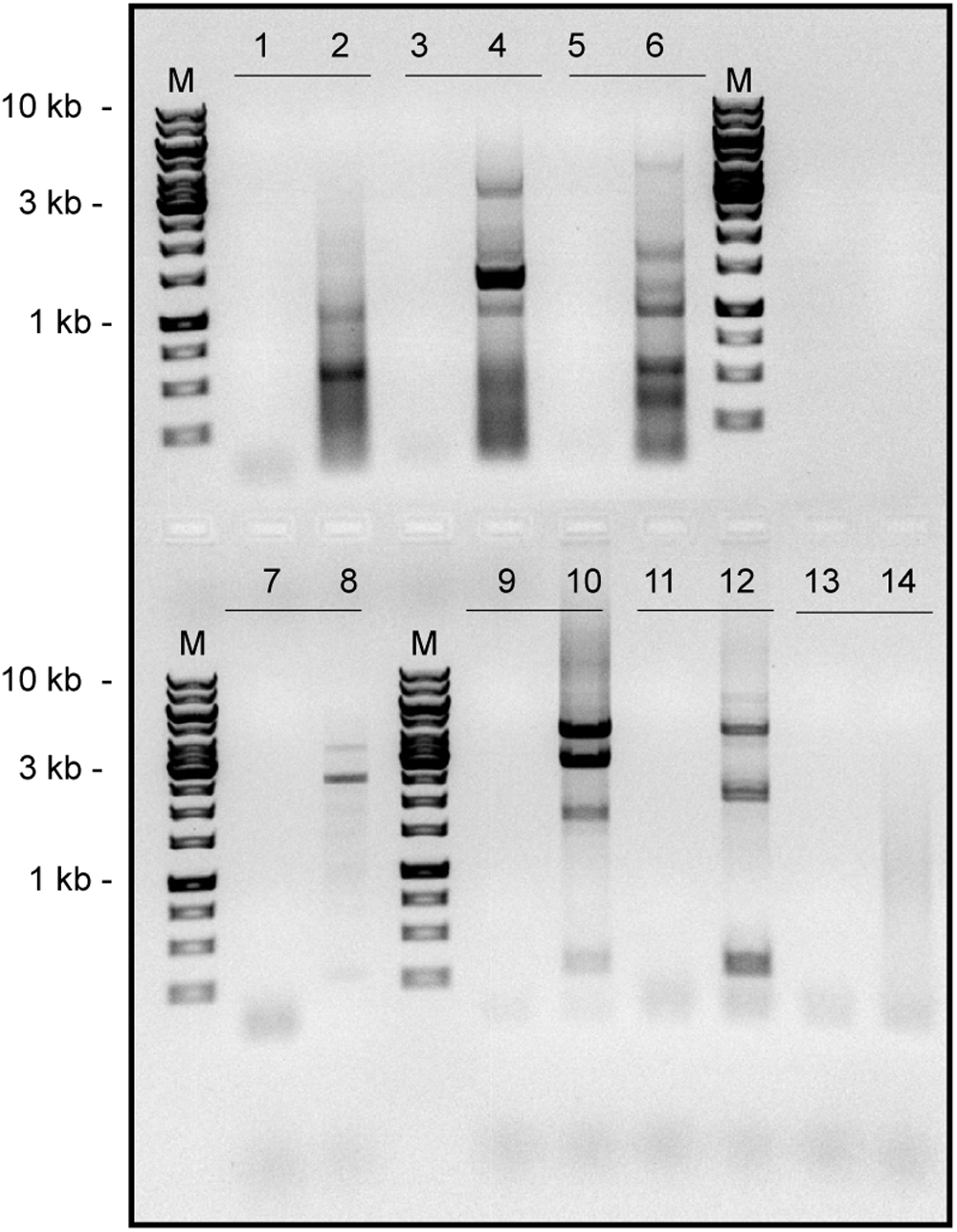
eccDNA iPCR. Inverse PCR products: lane 1, no template control (NTC) reaction; lane 2, G4 c.1 eccDNA; lane 3, NTC; lane 4, H3 eccDNA; lane 5, NTC; lane 6, G4 c.6 eccDNA; lane 7, NTC; lane 8, (ATTCT)47 eccDNA ; lane 9, NTC; lane 10, (CAG)102 c.10 eccDNA; lane 11, NTC ; lane 12, (CAG)102 c.13 eccDNA ; lane 13, NTC; lane 14, HeLa/406 DF/myc eccDNA (control, no microsatellite insert).

### Complex rearrangements in eccDNAs

The iPCR products from each microsatellite ES were analyzed by long read Hi-Fi circular consensus DNA sequencing (91,92). During the processing of the raw sequencing data, reads were deduplicated to eliminate sequence overrepresentation due to PCR amplification. Deduplication also eliminates all but one WT read, and any identical recombinants or mutants, i.e. all of the analyzed reads are unique.

The iPCR product sequences from the ectopic microsatellite cell lines were mapped by Ribbon (66), a tool which uses BWA-MEM alignment (60) against the ES sequence and the human reference genome (GRCh38 merged with the HeLa genome (27)) for the analysis of complex genome rearrangements. Due to the iPCR strategy, all reads contain domains that include the iPCR forward and reverse primer binding sites. The assignments in Ribbon were confirmed by the alignment of individual reads to the reference genome using BLAST (https://blast.ncbi.nlm.nih.gov). The Ribbon alignments stack individual reads from top to bottom by decreasing stringency scores, to plot the location of read sequences at the ES (left side of each panel) or to donor nonallelic chromosomes (right side). The *Reference Viewport* perspective in Ribbon (Figure 4) gives a broad picture of template switching within the ES and to nonallelic chromosomal sites in multiple reads.

**Figure 4.**
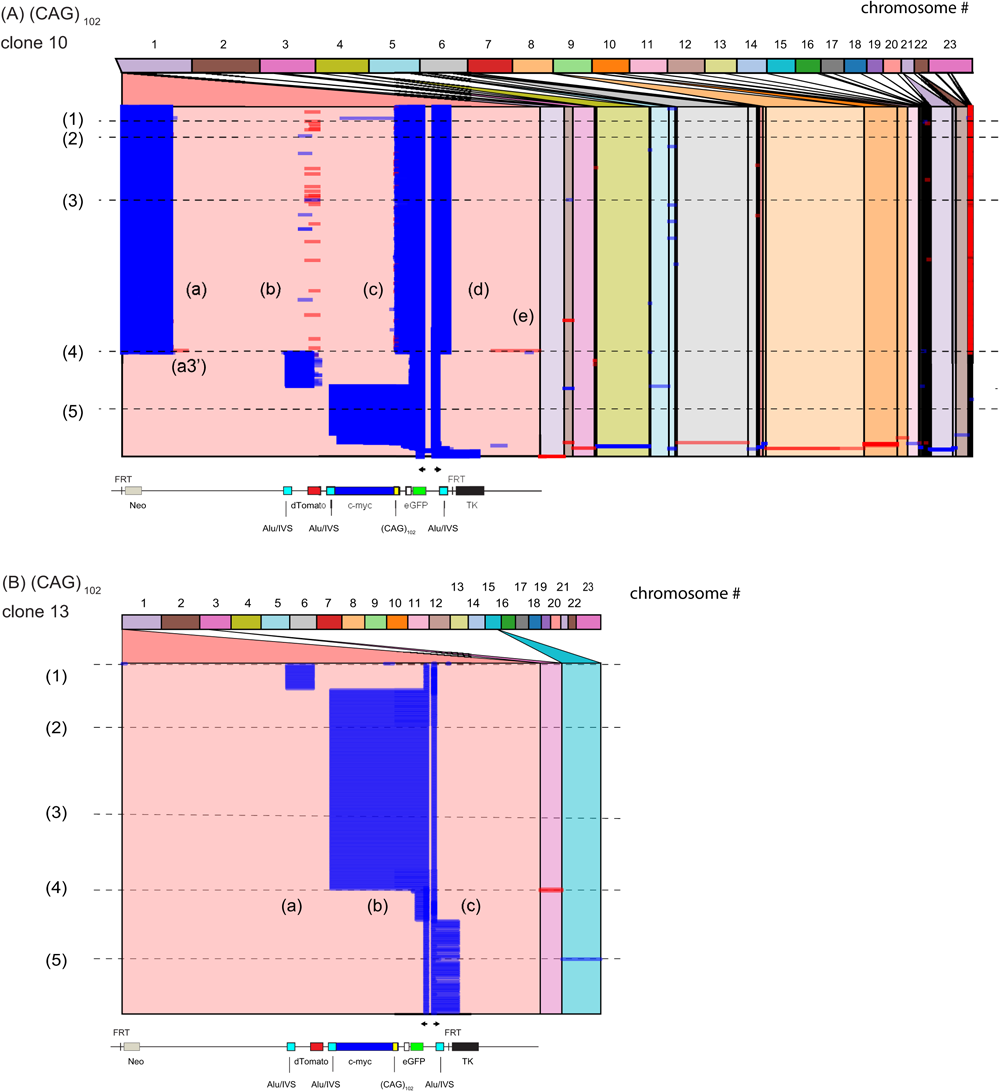
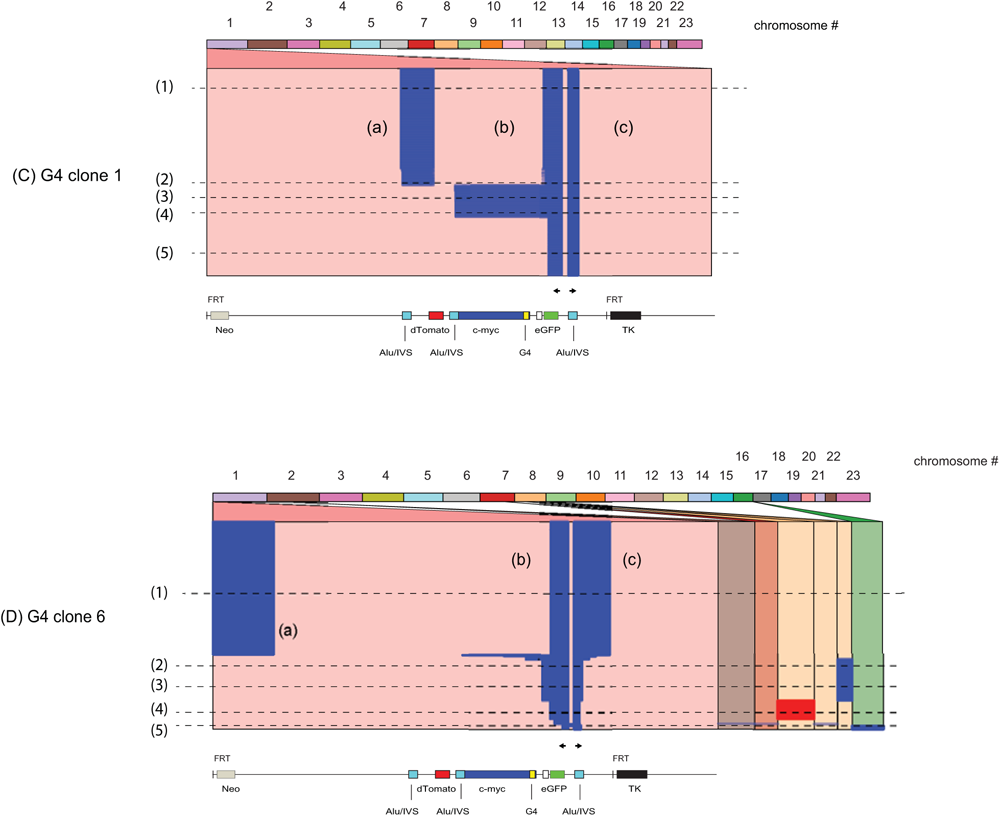
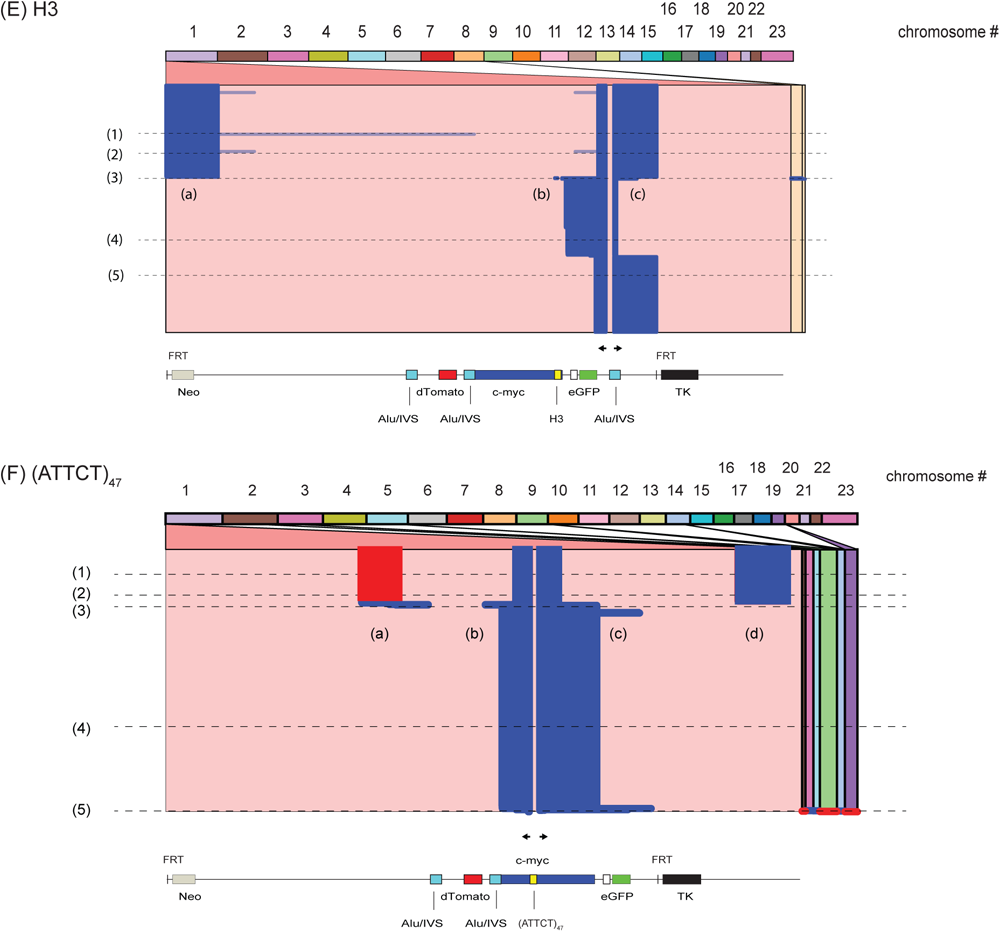
Template switching in eccDNAs. iPCR eccDNA reads were aligned using the Ribbon (66) reference viewport, and placed above a map of the ES; arrows indicate the approximate positions of the iPCR primers. (A) (CAG)102 c.10 eccDNA (n=26,466 reads); (B) (CAG)102 c.13 eccDNA (n=249 reads). (C) G4, c.1 eccDNA (n=485 reads); (D) G4 c.6 eccDNA (n=1695 reads). (E) H3 eccDNA (n=1197 reads); (F) (ATTCT)47 eccDNA (n=1299 reads). Cells were treated with siCON (Methods). Template switching domains (a), (b), (c) in panels (A)-(E) are labeled independenty. Blue, plus strand reads; red, minus strand reads. The ES region is located in the leftmost panel; nonallelic template switches are indicated in the righthand panels. Chromosome numbers are indicated across the top of each panel. Lines (1)-(5) in panels (A)-(E) are referred to in the text.

The most immediate observation from the reads obtained from any of the microsatellite cell lines is that large swathes of the ES are deleted from the eccDNAs. In addition, almost all (>97%) of the H3, G4 or (CAG)_102_ microsatellites were found to have been mutagenized in the eccDNAs. (The iPCR primers chosen for the (ATTCT)_47_ cell line did not report on the microsatellite sequence). Thus, eccDNAs undergo mutagenesis within the microsatellite and bidirectionally through proximal and distal flanking sequences.

We term those regions that are noncontiguous in the ES but are contiguous in the reads ‘template switching domains’. For example, the line 1 read in (CAG)_102_ c.10 cells (Figure 4A), shows five prominent template switching domains (a)-(e) within the ES that appear in the eccDNAs. It is notable that deletions due to template switching occur both upstream and downstream of the ectopic microsatellites and DSBs at the (CTG/CAG) and G4 non-B DNAs (22,77). These results likely exclude semiconservative mechanisms of eccDNA formation not involving DSBs, such as episome extrusion, due to differences in template switching between the nascent strands or nascent and template strands of a replication fork.

Strikingly, the same extended sequence (most often Alu or the 48 bp FRT), which we term an ‘eccDNA junction’ sequence, is present at the ultimate downstream ends of the domain(s) read by the forward primer and the upstream end of the domain(s) read by the reverse primer in 100% of the eccDNAs. This observation strongly suggests that homology directed repair is responsible for circularization of the eccDNAs.

For example, in (CAG)_102_ c.10 eccDNA reads above and including the line (4) read, the forward primer extension products (domain (d)) terminate at the downstream FRT sequence and recombine to form an eccDNA junction with the upstream FRT sequence at the 5’ end of domain (a). In contrast, several domain (d) reads below the line (4) read terminate at the downstream Alu d(T)_29_ repeat, and recombine to form eccDNA junctions with domain (c) products that terminate in the second Alu repeat or domain (b) products ending at the first Alu repeat.

Despite the extended junctions that circularize the eccDNAs, analysis of the internal template switching sequences shows that the template switching sites are not identical, but can vary by more than 250 bp (Supplementary Figure 1). Many of the (CAG)_102_ c.10 reads (e.g. Figure 4(A) line (3)) show template switches within the CAG repeats, while reads that cover the c-myc origin (e.g. line (5)) end at the d(T)_29_ sequence of the second Alu site, and consistently delete 30-60 CAG triplets.

The portion of the (CAG)_102_ c.10 eccDNA alignment below the line (4) read shows several reads in red with template switches to nonallelic chromosomes, indicating that these sequences were copied in the q (long arm) à p (short arm) direction.

In G4 c.1 cell reads above line (2), domain (a) becomes ligated to domain (b) at microhomologies between G4 consensus sites in the dTom and eGFP genes, deleting the entire c-myc region. Between lines (2) and (4) the c-myc origin is retained, but ∼250-260 bp extending from the intrinsic c-myc quadruplex forming sequences Pu27 (93) to the center of the G4 microsatellite insert is deleted in 100% of reads. In G4 c.6 cells and H3 cell reads above line (3) (Figure 4C), the circularization junction between domains (a) and (c) occurs through the 48 bp FRT homology. In contrast, the internal template switches to nonallelic sites on chromosomes 2, 7, 8 and 16 (read line (2) and below) occur through regions of Alu homology.

In (ATTCT)_47_ cells, the upper portion of Figure 4F shows that the nascent DNA strand reverses polarity to copy itself, or the parental strand of the sister template, in domain (a) (red reads), beginning at a 6 bp microhomology. In the reads below line (3) the nascent strand also copied domain (a), but in the forward polarity. Reads below line (3) show similar (+/-2 bp) internal template switching sites, and all show the same 8 bp circularization junction, located in the c-myc Pu27 (93) G4 consensus sequences. Several reads below line (3) also show 3’ extensions of domain (c) ending between eGFP and TK; all of these reads terminate within 24 bp of G4 consensus sequence matches. Taken together, these results suggest that quadruplex forming sequences are sites of heightened instability during the replication of eccDNAs.

The alignment pattern of (ATTCT)_47_ cells using iPCR primers flanking the ectopic microsatellite did not contain eccDNA sequences derived from the dTom gene (Figure 4F). However, when tested with iPCR primers flanking the dTom gene additional eccDNAs were revealed which contained dTom sequences (Supplementary Figure 2). Figure 4 also shows distinct eccDNAs that share only the iPCR primer domains. Taken together, these results indicate that distinct eccDNAs can be released from multiple regions of the same ectopic microsatellite site.

### Nonallelic template switches

Nonallelic chromosomal template switches were mapped based on BWA-MEM alignment using Circos (64), and are shown by lines connecting the ES to nonhomologous donor chromosomes (Supplementary Figure 3). Each of the microsatellite cell lines, except for G4 c.1, showed nonallelic chromosomal template switches. Chromosome 8 was a target for template switching in several of the cell lines. Since chromosome 8 harbors the c-myc gene, and the 5’ portion of that gene is part of the ES, we wished to confirm that the template switches are not BWA-MEM misalignments. BLAST sequence analysis showed that 34 of 357 reads from (CAG)_102_ c.10, 2 of 3 reads from (CAG)_102_ c.13, all 159 of the reads from G4 c.6, and 12 of 17 reads from H3 cells contain sequences on chromosome 8 that do not appear at the ES. Similarly, the PGK promoter from chromosome X (chromosome 23) is used as a promoter for the dTom gene; (CAG)_102_ c.10 shows two template switches to chromosome 23, neither of which is at the PGK locus.

At the resolution of these Circos plots, apparently identical breakpoint junctions may be many kilobases apart on the nonallelic chromosome. Thus, (CAG)_102_ c.13 template switches to chromosome 8 are separated by ∼49 kb, while G4 c.6 exhibits 13 template switches to the long arm of chromosome 8 that are ∼480 kb apart. This analysis reinforces the view that template switching during the formation of eccDNAs leads to recombination between sequences within the ES and to nonallelic chromosomes.

### eccDNA Hypermutation

We observe an apparent rate of mutation (indels, base substitutions) in the mapped eccDNAs of approximately 1.2 x 10^-2^ mutations/kb/generation. However, recent analyses of long read synthesis-based sequencing have reported that polymerase pausing at potential non-B DNAs (direct/inverted/mirror repeats, G4 motifs, A-phased repeats) can decrease or increase the apparent mutation rate in PacBio raw sequence reads by -1.23 fold (G4 motif complement) to +1.79 fold (G4 motif) in the non-B DNA sequences relative to non-motif regions (frequency ∼0.2 mutations/kb) (94). These effects are reduced ∼3 fold by circular consensus sequencing (ccs) (94–97), as in this work.

To test whether this issue affected our conclusions, we confirmed the error frequency of PacBio ccs sequencing in the same ES DNA (non-microsatellite) of four plasmids used to construct non-B DNA clones (<0.2 errors/kb). It has also been suggested that slow-down of the replicative polymerases in vivo at the same structures as sequencing polymerases in vitro contribute to in vivo mutagenesis (94). To assess the possible contribution of errors during sequencing, we compared the range of mutation rates of the clones tested here after removing reads below a threshold of 2 mutations/kb from consideration. The high mutation threshold resulted in a range of mutation rates of 1.1 x 10^-2^ mutations/kb/generation [in (ATTCT)_47_ cells] to 1.8 x 10^-2^ mutations/kb/generation [in (CAG)_102_ c.13 cells] compared to the range of rates without applying a mutation threshold of 1.1 x 10^-2^ mutations/kb/generation [in (CAG)_102_ c.13 cells] – 1.4 x 10^-2^ mutations/kb/generation [in G4 c.1 cells]. The high mutation threshold did not appreciably change the appearance of the alignments of the (CAG)_102_ clones (Supplementary Figure 4), the G4, H3, or (ATTCT)_47_ cell lines (not shown), or the mechanistic conclusions of this work. Therefore, no threshold was subsequently applied to the data.

The *Query Viewport* perspective in Ribbon shows the arrangements of the template switching domains of individual eccDNA reads. These plots (Figure 5, Supplementary Figure 5) show that both ES sequences and nonallelic template switch domains display hypermutagenesis, and that reads containing the same switching domains display distinct patterns of mutagenesis.

**Figure 5.**
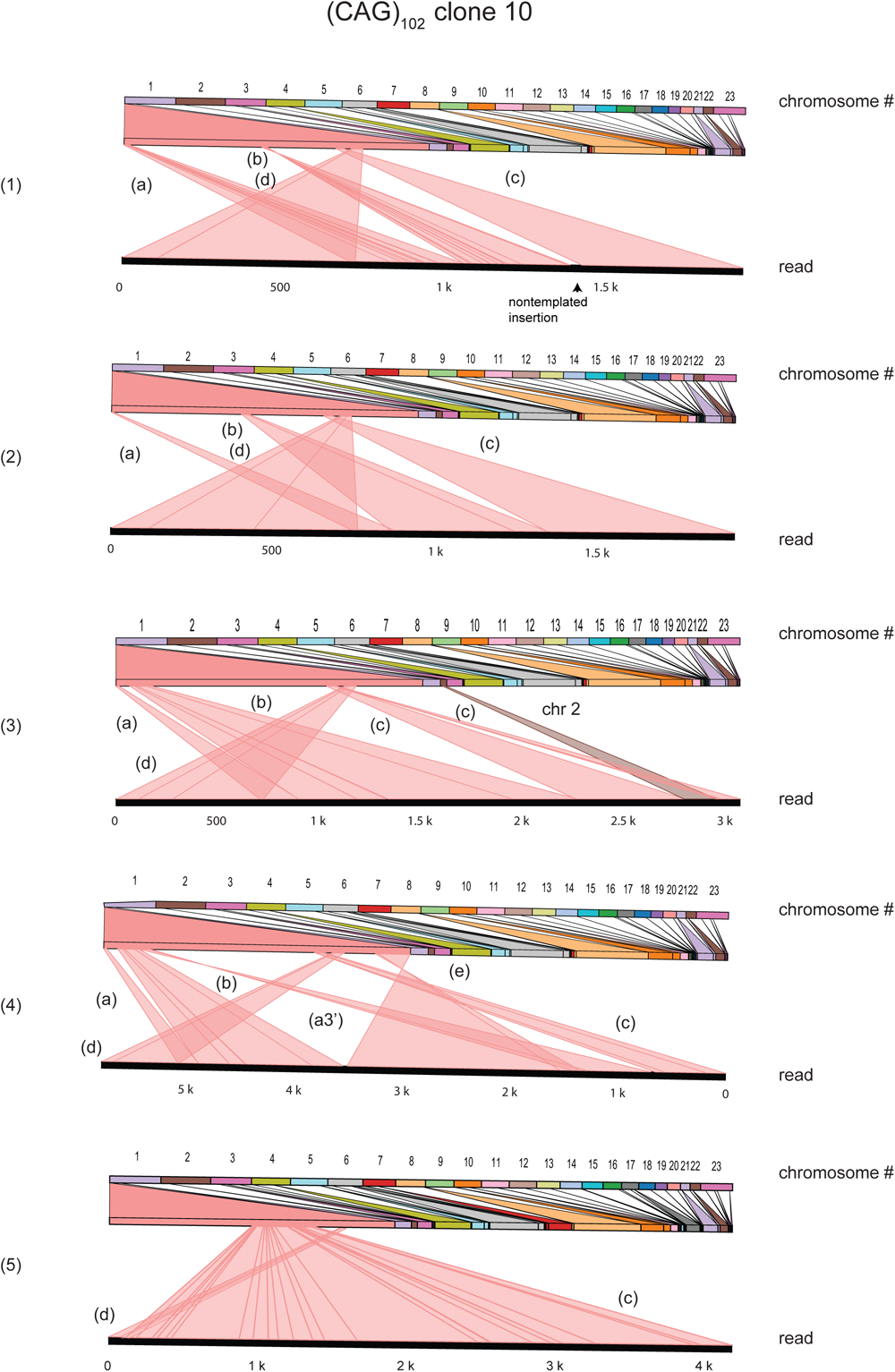
Template switching patterns of eccDNAs in (CAG)102 clone 10. Individual reads from (CAG)102 c.10 cells (lines (1)-(5) Figure 4A) were mapped in the query viewport perspective by Ribbon. Chromosome numbers are shown at the top; the lower heavy black line in each panel is the complete read. Heavier red lines within a domain represent indels. Letters (a), (b), (c), (d) correspond to template switch domains in Figure 4A.

The query viewport views of (CAG)_102_ c.10 read lines (1) - (5) are shown in Figure 5. Clone (CAG)_102_ c.10 line (1), (2) and (3) reads encompass domains (a), (b), (c) [the reverse primer domain] and (d) [the forward primer domain], but show different patterns of indels and substitution mutations. A schematic diagram (Figure 6) shows the structure of the (CAG)_102_ c.10 line (1) read, the corresponding eccDNA, and the FRT junction homology between domains (a) and (d). Reads below line (3) show similar (+/- 2 bp) internal template switching sites, but large differences in indels (+/- 200 bp) and base substitutions.

**Figure 6.**
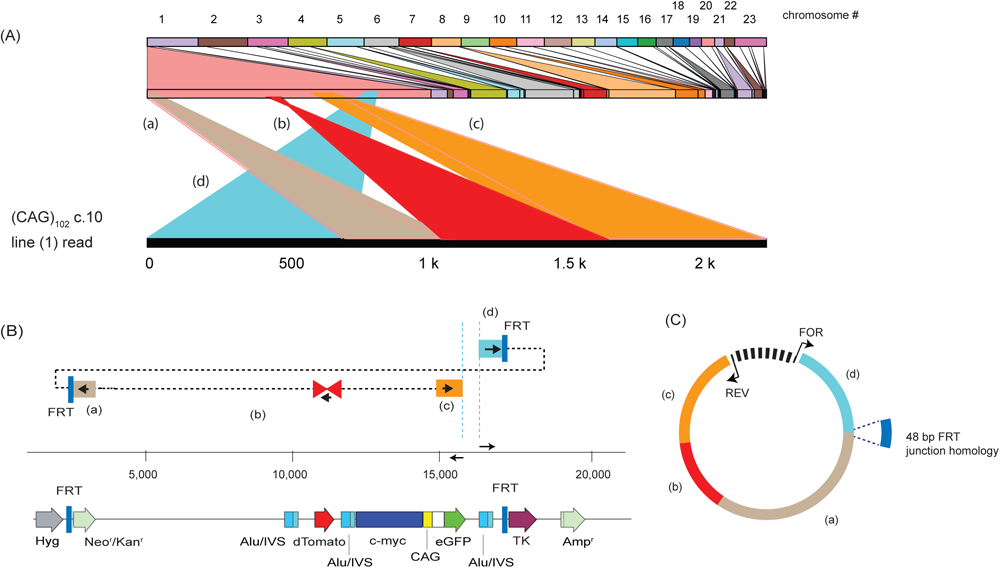
Template switching schematic of (CAG)102 clone 10, line 1 read. (A) Query viewport view (Figure 4A, line 1) of template switch domains (a)-(e). (B) Interpretation of template switches aligned with the ES map. Arrowheads within the domains indicate the direction of BIR strand extension. (C) Circular map of the eccDNA derived from the (CAG)102 clone 10, line 1 read.

Overall, distinct overall mutagenesis patterns of the H3, G4 and (CAG)_102_ microsatellites are evident in the query viewport views. In outline, all H3 reads are mutagenized at the triplex forming sequence; mutations are base substitutions, internal indels, and insertions at the 3’ microsatellite boundary. In contrast, the quadruplex-forming sequences are deleted completely in G4 c.1 and c.6 cells, while (CAG)_102_ clones show primarily short (5-150 bp) deletions within the microsatellite, or insertion of incomplete CAG repeats at 3’ edge of the microsatellite.

The mutations in different eccDNAs could arise independently or progressively in a cell lineage as a chromosomal ES accumulates successive changes. Given the likelihood of autonomous eccDNA replication (98,99), different mutational patterns might also arise consecutively in an eccDNA lineage. Thus, multiple G4 c.1 domain (b) reads (Supplementary Figure 5B) show identical template switch sites but different sets of indels and base substitutions. For example, G4 c.1 line (3) and (4) reads across domains (b) and (c) have the same template switch junctions but substantially different mutational patterns. Similarly, G4 c.6 reads show identical template switch sites with different sets of mutations. In contrast, G4 c.1 line (2), (3), and (5) reads contain different nonallelic template switches, and G4 c.6 cells show distinct nonallelic template switches to chromosome 8 (read lines (3) and (4)) in the p à q direction and in the q à p direction, respectively.

In H3 cells, the line (3) read crosses the H3 microsatellite but mutates 28 bp of 60 bp of the H3 microsatellite sequence, while the 5’ ends of domain (b) between the line (3) and (4) reads begin ∼200 bp downstream from the H3 microsatellite. In these cells, the line (1) read is the longest (>10 kb) read analyzed and exhibits a high mutation frequency (∼1.33 x 10^-2^/bp, ∼5x10^-5^ mutations/bp/generation) despite the read not covering the microsatellite domain, consistent with mutagenesis at distances greater than five kilobases from the ES. In the H3 line (4) read, the domain (b) and (e) sequences are criss-crossed (minus strand reads), indicating that the replicative polymerase had reversed direction and switched templates to copy the upstream (self) nascent DNA, or the sister chromatid parental DNA.

Insertions that are not found in the human, bacterial or viral NCBI GenBank databases are present at apparent gaps in the sequences of H3 cell reads (3) and (4), (ATTCT)_47_ read 3, and G4 reads (Supplementary Figure 5). These segments may be due to mutagenic misalignment insertions by the TLS POLθ (100–104), which has been implicated in off-target integration events during DSB repair (104–106).

In (ATTCT)_47_ cells, the nascent strand reversed polarity to copy itself or the parental strand sequences of the sister chromatid in the reads of domain (a) (Figure 4C, red). The line (1) and line (2) reads show similar template switching domains, however, these reads differ dramatically from one another in the extent and nonoverlapping patterns of mutagenesis, and in distinct template switch boundaries (Supplementary Figure 5), suggesting that these rearrangements occurred in independent progenitors, rather than as successive events in a single lineage.

Altogether, the query viewpoint patterns show that (a) indel and base substitution mutagenesis occurs upstream and downstream of each microsatellite, in both ES and nonallelic sequences, (b) multiple indel and substitution mutations are often observed at local hotspots, and (c) reads with similar template switches display independent patterns of mutagenesis over the same DNA sequences.

### Microhomology-dependent template switching

Short sequence homologies (<70 bp) are often found to overlap breakpoint junctions repaired by nonhomologous end joining (NHEJ), ALT-NHEJ (microhomology-mediated end joining, MMEJ), ‘fork stalling and template switching’ (FoSTeS) and MMBIR (107–115). Short sequence homologies also occur alongside break-point junctions without frankly overlapping the junctions (116–119), in agreement with the notion that sequences upstream of a strand break play a role in determining template switching junctions.

Reads from (CAG)_102_ c.10 were analyzed by ALVIS (63), a tool for the visualization of contig overlaps (Figure 7). The template switching domains show overlaps of 0 - 48 bp of homology. The base position of the overlaps varies between reads, although several reads show similar patterns in addition to common domain boundaries. Consistent with the contribution of Alu elements to genomic plasticity (29,120–122), a distinct hotspot for template switching from the second to the third Alu element (Figure 7 (A)-(E), (G), (H)) is found within the Alu d(T)_29_ tail. The same sequence is a preferred site for ES and nonallelic template switching in (CAG)_102_ c.13 cells (Supplementary Figure 6A) where overlap homologies as long as 121 bp were observed. In (CAG)_102_ c.10 and c.13 cells, template switching within the Alu d(T)_29_ sequence consistently resulted in the deletion of 2-4 d(T) nucleotides.

**Figure 7.**
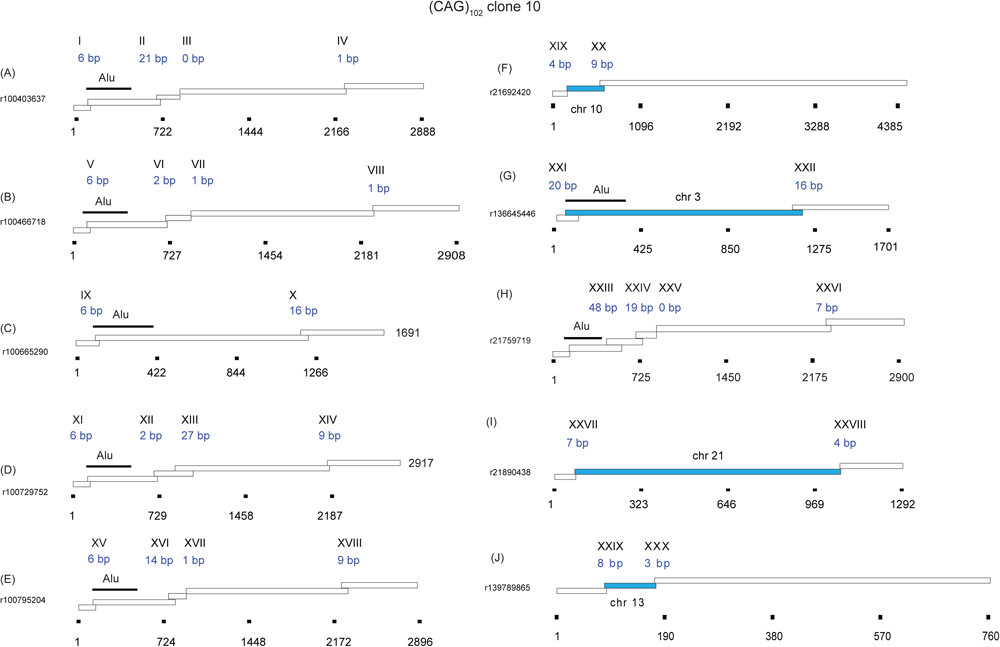
Microhomology overlaps in (CAG)102 clone 10. (a)-(j) are individual eccDNA reads from (CAG)102 clone 10 cells. Each box is a template switch domain. Blue domains are nonallelic template switches. Roman numerals and blue base pair numerals are sequence overlaps. Approximate length coordinates are shown below (or alongside) each read. Analysis was performed with ALVIS (178) and reformatted for presentation.

Clones containing the G quadruplex-forming consensus sequence, G4 c.1 and G4 c.6 also show frequent recombination at the d(T)_29_ sequence (Supplementary Figure 6B, C), however, the overlap between Alu sites is markedly shorter in the ES recombinants than in the template switches between chromosomes. A pattern of 6 bp overlap in ES template switches between Alu elements is seen in G4 c.1 and G4 c.6 reads, whereas nonallelic template jumps, particularly to chromosome 8, displayed 43-101 bp overlaps. Thus, different mechanisms may be involved in intrachromosomal vs. nonallelic template switching, potentially reflecting disparate roles of Rad51 in these processes (54,123).

The G4 c.1 and G4 c.6 reads also show several insertions of nonrepetitive, nontemplated insertions of 30-60 bp, consistent with the proposed activity of DNA polymerase θ in MMBIR and theta-mediated end joining (TMEJ) (101,102,119). We note also that the chromosome 8 template switches recurrently target Alu sequences 27 kb upstream of the c-myc locus. Supplementary Figure 6 also shows several reads from G4 c.6 with template switches flanking, but not overlapping, an Alu repeat. This is in line with the idea that homeologous sequences upstream of the invading 3’ end are involved in discriminating between multiple potential microhomologous sites for invasion (115,117).

Template switches in H3 cells also frequently occur with 6 bp overlaps at the beginning of the Alu d(T)_29_ sequence (Supplementary Figure 6D), however, a fraction of recurrent template switches to chromosome 8 do not occur within Alu repeats, and do not target the c-myc locus, but show template jumps 1.5 Mb upstream of c-myc. The differences in the patterns of nonallelic template switches in the G4 vs. H3 clones suggest that these non-B DNAs may differ in the way they remodel the stalled replication fork (124). Consistent with this suggestion, (ATTCT)_47_ cell reads rarely show template switches within Alu repeats, and those reads that show Alu recombination (e.g. Supplementary Figure 6E) do not recombine within the d(T)_29_ Alu sequence.

We also wished to investigate whether there is regularity or randomness to the pattern of microhomology selection. As shown in Figure 8, there is a clear preference for 10 bp overlaps in G4 c.1 cells, but a dominance of 1 bp, 2 bp, 6 bp, and 8 bp overlaps in G4 c.6 cells, and overlaps of 2 bp and 5 bp in clone H3. Clone (ATTCT)_47_ reads show a majority of 2 bp, 4 bp, 6 bp and 22 bp overlaps. In contrast, reads from clone (CAG)_102_ c.13 show a preponderance of 14 bp overlaps, while clone (CAG)_102_ c.10 shows overlaps of 1 bp, 2 bp, 8 bp, 9 bp, 13 bp, 16 bp and 42 bp. When challenged against a random number array, each of these distributions is highly nonrandom. We suggest that sequences preceding the 3’ end contribute to the selection against multiple potential microhomology targets, and that the nonrandom patterns may reflect the activities of distinct DNA damage tolerance polymerases.

**Figure 8.**
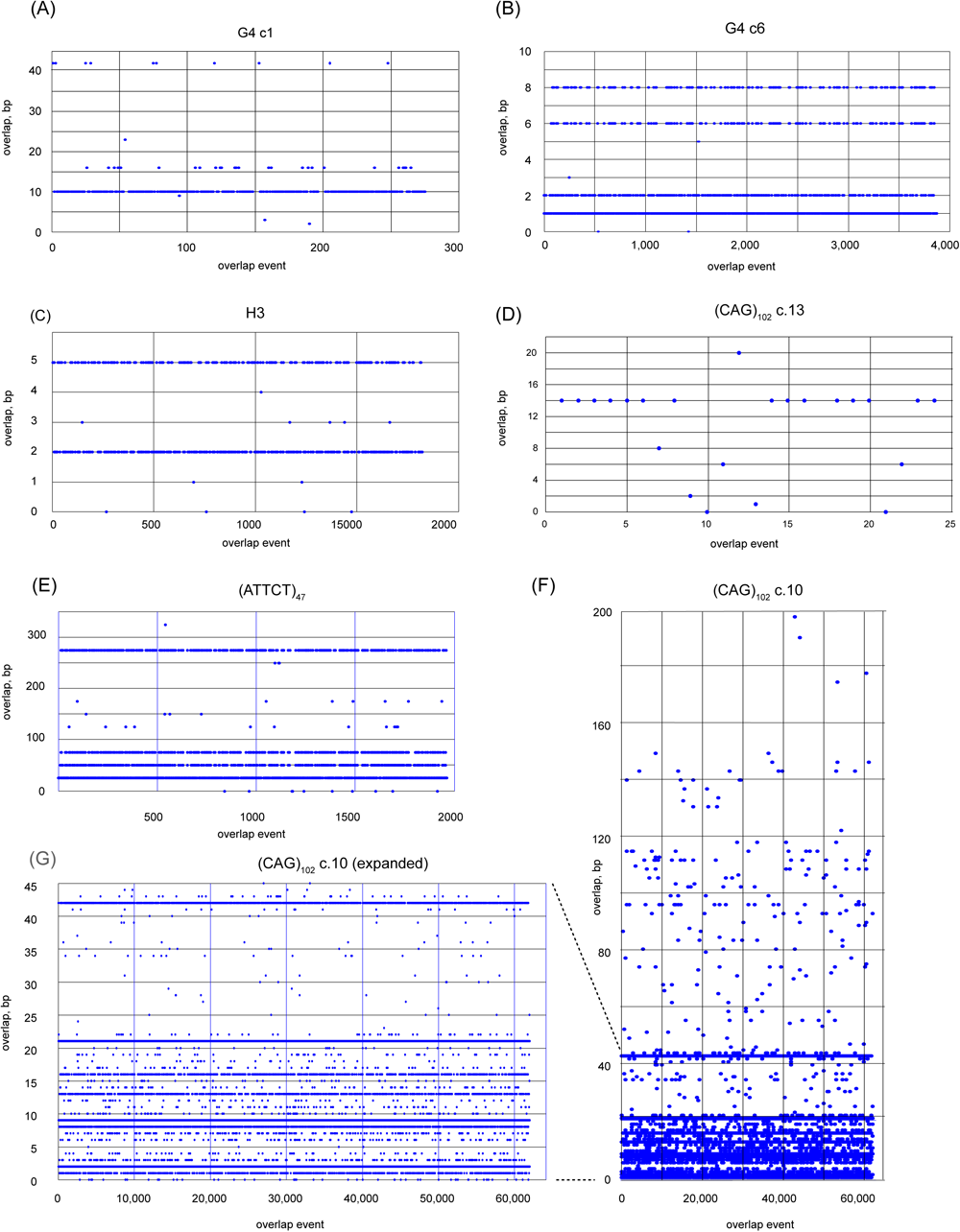
Overlap microhomology. Distribution of sequence overlap homology between eccDNA template switching domains in (A) G4 clone 1 cells; (B) G4 clone 6 cells; (C) H3 cells; (D) (CAG)102 clone 13 cells; (E) (ATTCT)47 cells; (F, G) (CAG)102 clone 10 cells. Y-axis, overlap base pairs; X-axis, overlap event number. Note that a read with several template switches will have multiple overlap events. Each of these distributions is nonrandom and distinct from one another (p<10-30, pairwise Student’s t-tests).

### Single molecule analysis of mutations

The approximate location of mutations in the numbered reads of Figures 4 were mapped in BLAST by alignment to the ES or to the reference GRCh38/HeLa genome, and are shown in Figure 9 and in Supplementary Figure 8. The frequency of mutations (indels, substitutions) in these reads ranged from ca. 2 x 10^-3^/bp to 1 x 10^-2^/bp.

**Figure 9.**
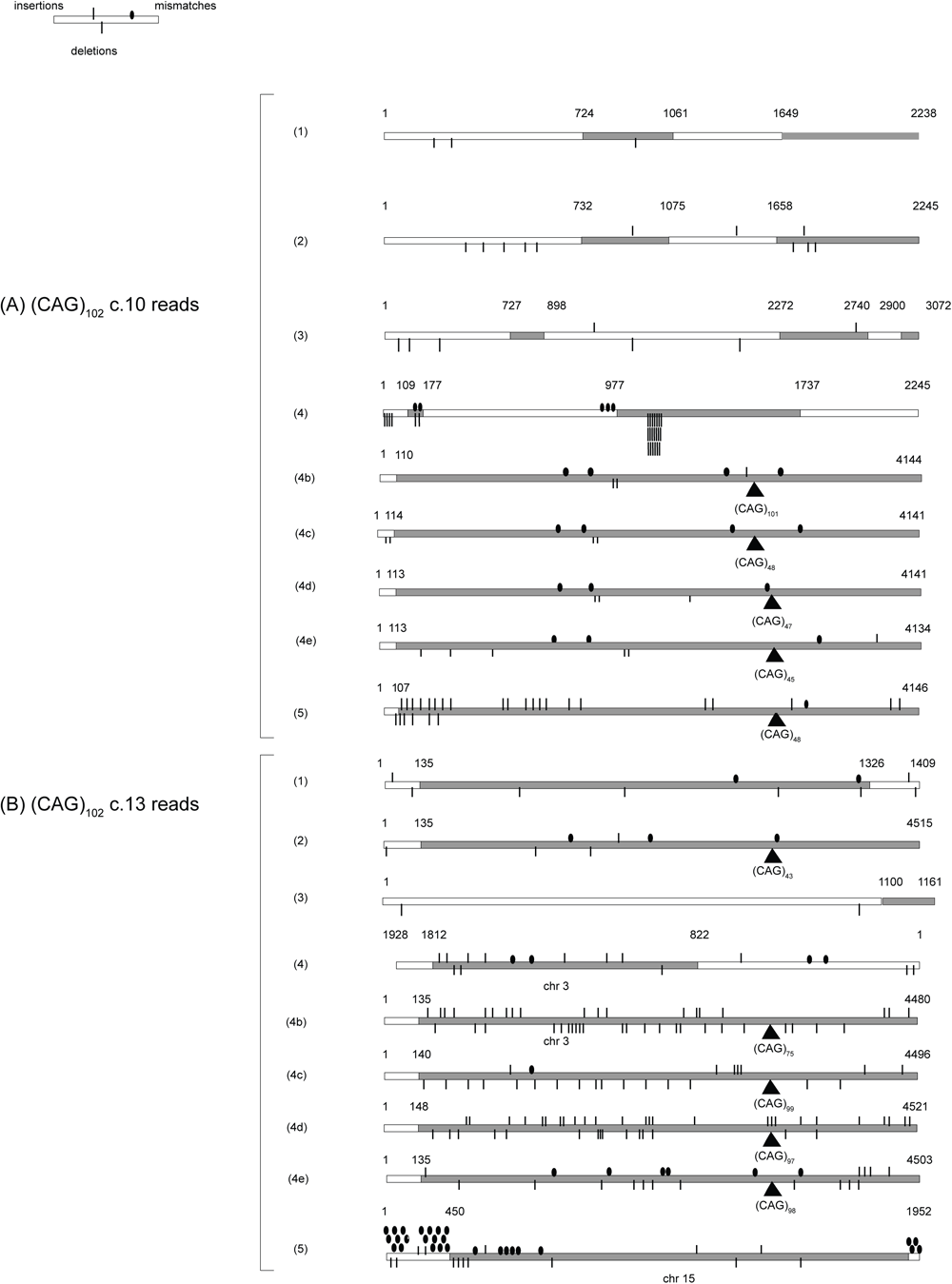
Mutation patterns of (CAG)102 eccDNAs. The approximate distribution of mutations mapped in BLAST for (A) (CAG)102 c.10, read lines (1)-(5); (B) (CAG)102 c.13 read lines (1)-(5). Lines (4b, 4c, 4d, 4e) are additional examples of neighboring reads between lines (4) and (5) of (CAG)102 c.10 and c.13 (from Figure 4).

We previously mapped a MUS81-dependent DSB immediately 3’ to the ectopic (CTG)_102_ microsatellite and showed that hypermutagenesis occurred 5’ and 3’ from that DSB (22). The spread of mutagenesis upstream and downstream from the hairpins in (CAG)_102_ c.10 cells and (CAG)_102_ c.13 cells (59) shows further evidence of replication-dependent mutagenesis, consistent with a BIR-like repair mechanism extending bidirectionally from a double-ended DSB. One of the longest reads (Supplementary Figure 8; H3, line 1) shows that hypermutagenesis extends more than 10 kb upstream from the H3 microsatellite repeat. Additionally, reads covering the same region of the ES (e.g. H3 cells, lines (2), (2B); (3), (3B); (ATTCT)_47_ cells, lines (1), (1B); (2), (2B);) show dramatically different frequencies and positions of mutations, suggesting that the activity of the repair replisome over the same template sequence differs between repair events (21).

We analyzed the mismatch mutational signatures of the eccDNAs in the absence of external replication stress, and of the (CAG)_102_ and (ATTCT)_47_ clones in the presence of drug-or protein knockdown-induced replication stress (Supplementary Figure 8). Each bar represents the sum of the substitutions for the central base of the sixteen possible trinucleotides. The high levels of C/G>T/A mutations seen in H3, G4 and ATTCT clones are characteristic of APOBEC cytidine deamination, while the elevated T/A>G/C single base substitution mutations seen in (CAG)_102_ clones are characteristic of POLη mutagenesis (125). Overall, the signatures do not identify the mutation signature of any single TLS polymerase but resemble COSMIC (https://www.sanger.ac.uk/tool/cosmic/) Signature 3 (119), ascribed to homologous recombination repair deficiency (119). The mutation frequency of these cell lines varied between ∼0.2-0.4 single base substitution mutations/kb except for a larger increase in the mutation rate and C/G>A/T conversion of G4 clone 1, as noted).

### Replication stress-dependent changes in eccDNA

To test the replication-dependence of eccDNA formation, we assessed whether low doses of the deoxyribonucleotide synthesis inhibitor hydroxyurea (HU, 0.2 mM) or the replicative polymerase inhibitor aphidicolin (APH, 0.2 uM) would alter the structure of eccDNAs, based on previous results showing that these treatments resulted in replication stress and enhanced microsatellite instability (59). Treatment of (CAG)_102_ c.10 cells with HU resulted in the dramatic decrease of template switching, including the loss of template switching domains (a) (b) and (e), and the loss of reads that extended from the c-myc origin to nonallelic chromosomal sites (Figure 10). Treatment of (CAG)_102_ c.10 cells with APH resulted in the loss of reversed (red) reads potentially stemming from fork regression, and the narrowing of domain (d) reads. These replication inhibitors also changed the eccDNA template switching patterns of (CAG)_102_ c.13 cells where HU treatment severely limited eccDNA to the region around the iPCR primers, while APH treatment resulted in the loss of reads containing domain (b) and reduced nonallelic recombination. In addition, treatment with HU or APH, changed the base substitution signatures of both (CAG)_102_ clones, consistent with the recruitment of alternative error-prone polymerases to stalled forks under replication stress. The query viewport patterns for individual read lines (1) - (5) of (CAG)_102_ cells treated with HU or APH are shown in Supplementary Figure 7.

**Figure 10:**
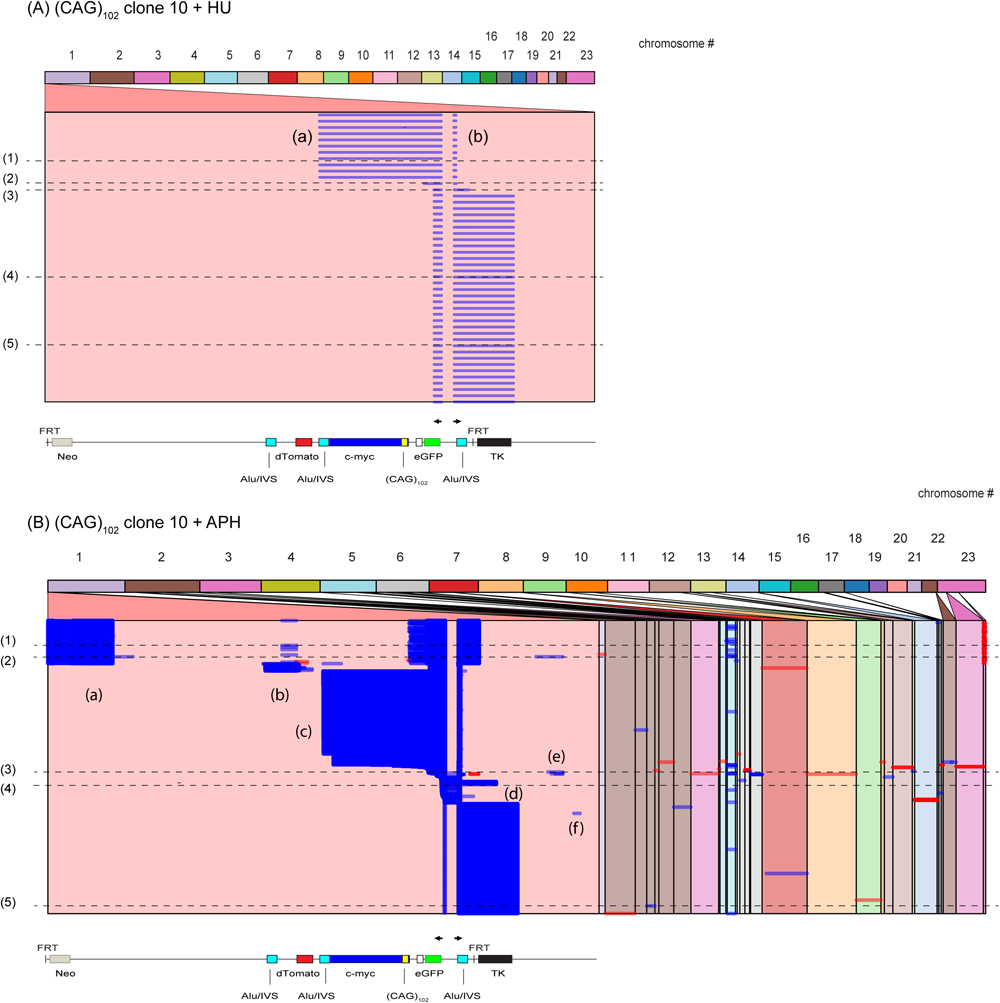
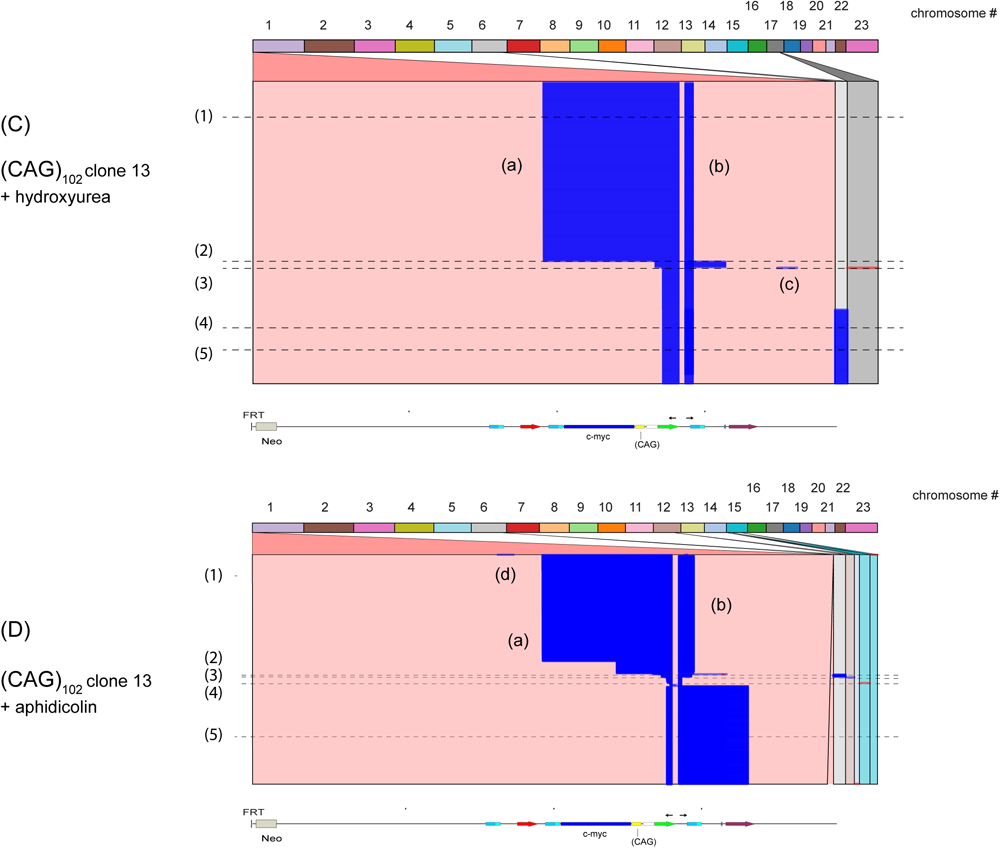
Effects of replication stress on eccDNA structure. Alignment of iPCR reads from (CAG)102 c.10 cells treated with (A) hydroxyurea (n=18 reads), or (B) aphidicolin (n=2439 reads). Alignment of iPCR reads from (CAG)102 c.13 cells treated with (C) hydroxyurea (n=264 reads), or (D) aphidicolin (n=736 reads).

The increased instability of the CAG microsatellite ES’s by drug treatment is consistent with the induction of replication stress (7,29,126–129) and the effects of HU and APH on expanded CTG microsatellites (22). We conclude that perturbation of replication alters the patterns of template switching in eccDNAs, likely due to remodeling of stalled replication forks. We note as well that HU and APH each have different effects on eccDNA structure, indicating that alternative forms of replication stress have different consequences for microsatellite instability (86,87), possibly due to the alternative folding of stalled forks. Because HU or APH were administered only for 48-96 hours before DNA isolation, the changes in eccDNA structure that were induced in (CAG)_102_ cells by these treatments are recent compared to the structures that accumulated during outgrowth of the clonal cell lines. Therefore, mutagenesis is ongoing in these cell lines.

### Rad51 affects eccDNA template switching

Rad51-dependent and -independent forms of BIR have been described in S. cerevisiae (130–133). To test whether Rad51 plays a role in the homology search leading to nonallelic template switching, or in the reversal of stalled forks (54,97,134–140) during eccDNA synthesis, we knocked down Rad51 in (CAG)_102_ c.10 and (CAG)_102_ c.13 cells with shRNA (Figure 11). Rad51 depletion resulted in dramatic changes in the pattern of reads in (CAG)_102_ c.10 cells (decreased domain (a) and (b) reads, loss of reversed reads in domain (b)) and in (CAG)_102_ c.13 cells (decreased domain (a) and (b) reads, increase of domain (c) read length).

**Figure 11:**
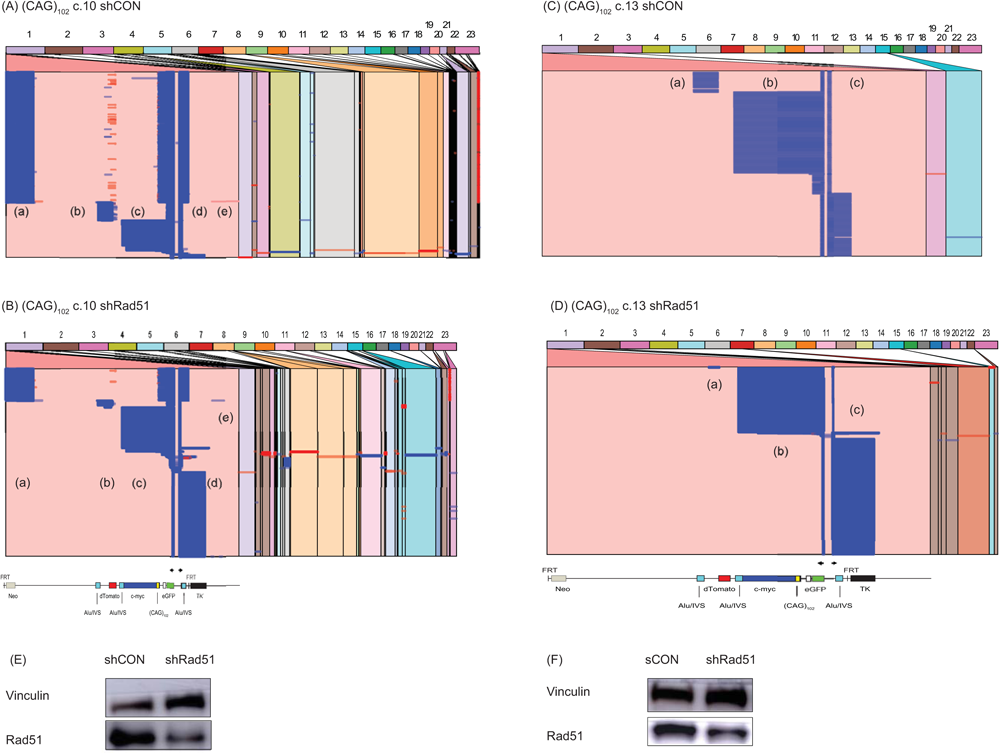
Remodeling of replication forks following Rad51 depletion. Alignment of iPCR reads from (A) (CAG)102 clone 10 cells, (B) (CAG)102 clone 10 cells depleted of Rad51. (C) Western blot (CAG)102 clone 10 cells. (D) Alignment of iPCR reads from (CAG)102 clone 13 cells, (E) (CAG)102 clone 13 cells depleted of Rad51. (F) Western blot, (CAG)102 clone 13 cells.

Comparing the lengths of template switch overlaps in control vs. Rad51 depleted (CAG)_102_ cells, we observed significantly decreased overlap lengths for both sister chromatid (or self) template switches, and for nonallelic template switches in Rad51 knocked down cells (Figure 12). In addition, depletion of Rad51 changed the base substitution signatures of both (CAG)_102_ clones (Supplementary Figure 8). In agreement with evidence for multiple roles of Rad51 in fork reversal (54,97,137,139–141) and recombination (142–144), our data illustrate pleiotropic functions for Rad51 in the remodeling of stalled forks and in homology searches during template switching in Rad51-dependent BIR.

**Figure 12:**
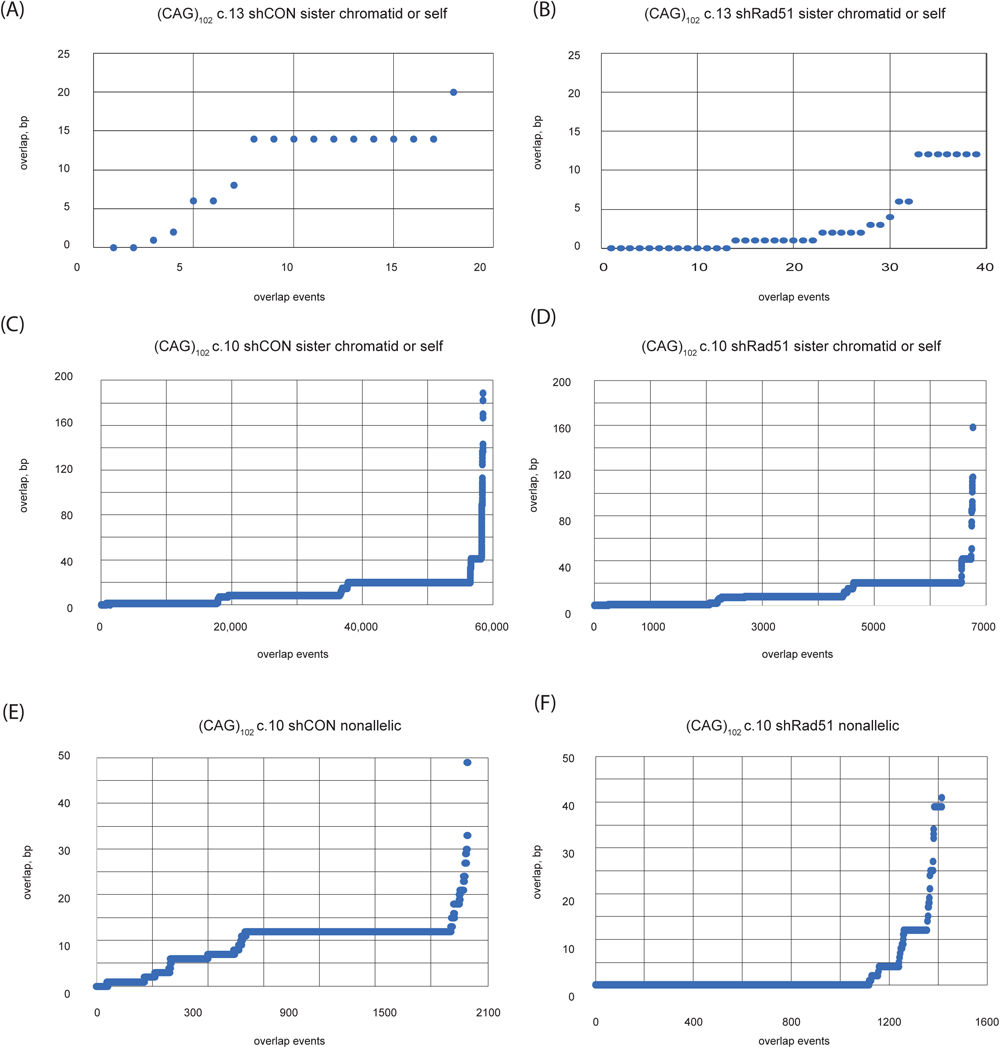
Effect of Rad51 depletion on template switching overlaps. Overlap homology of template switching to the sister chromatid or self (nascent) DNA in (A) (CAG)102 c.13 cells, (B) (CAG)102 c.13 cells depleted of Rad51. (C) (CAG)102 c.10 cells, (D) (CAG)102 c.10 cells depleted of Rad51. Overlaps of template switching to nonallelic sites in (E) (CAG)102 c.10 cells, (F) (CAG)102 c.10 cells depleted of Rad51. P values: (A) vs. (B), p= 9.74x10-6; (C) vs. (D), p = 7.88x10-53; (E) vs. (F), p = 7.77x10-6. (Student’s t-test).

**Figure 13:**
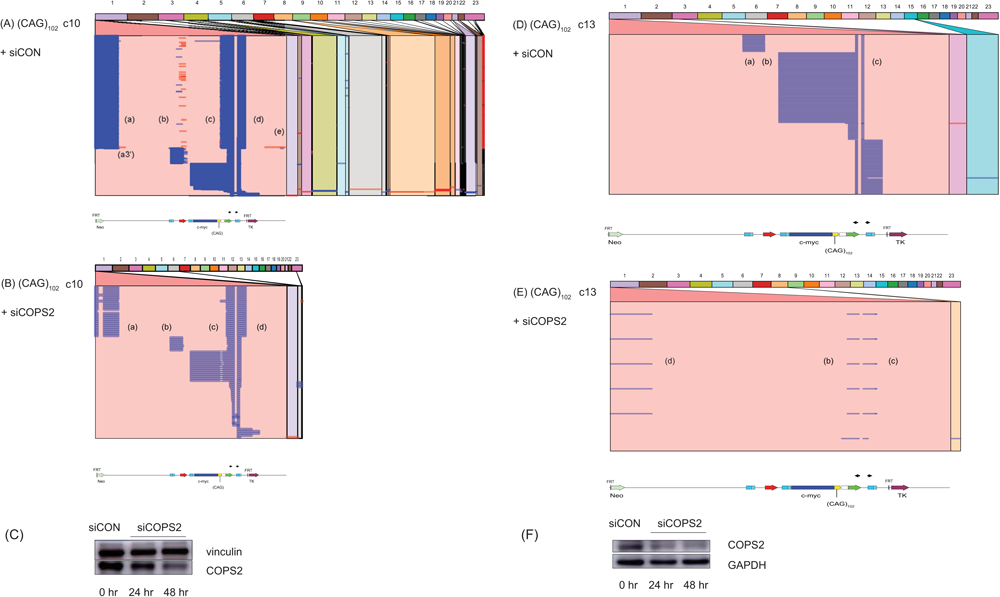
Effect of COPS2 depletion on eccDNA structure. Alignment of iPCR reads from (A) (CAG)102 clone 10 cells, (B) (CAG)102 clone 10 cells depleted of COPS2. (C) Western blot (CAG)102 clone 10 cells. (D) (CAG)102 clone 13 cells, (E) (CAG)102 clone 13 cells depleted of COPS2. (F) Western blot, (CAG)102 clone 13 cells.

### Multiple effects of COPS2 and TLS on eccDNA structure

We recently reported the results of an shRNA screen for suppressors of BIR mutagenesis, which identified the COPS2 subunit of the COP9 signalosome (20). COP9 inactivates the CUL4A/B ubiquitin ligase that targets PCNA (145), and thereby inhibits TLS polymerase POLη binding and TLS polymerase switching (146–151). Here, knockdown of COPS2 resulted in a decrease of eccDNA reads from an equivalent amount of total genomic DNA (Figure 12), and in substantial changes in the structure of eccDNAs in (CAG)_102_ c.10 cells (decreased nonallelic template switches, loss of template switching (reverse reads) in domain (b), introduction of an additional template switch within domain (a), deletion of ∼45 CAG repeats) and in (CAG)_102_ c.13 cells (loss of domain (a) reads, appearance of domain (d) reads, shortening of domain (b) reads). We therefore wished to examine whether the effects of COPS2 depletion could be recapitulated by knockdown of the POLη TLS polymerase.

TLS polymerases have been implicated in microhomology-mediated BIR (MMBIR) in yeast (112), and we have reported that knockdown of Rad18, POLη or POLκ increases instability at an ectopic (CTG)_100_ microsatellite (22). To determine whether the loss of POLη activity could account for the effects of COPS2 knockdown, we tested the effect of POLη knockdown on eccDNA structure. As shown in Figure 14, POLη depletion had little effect on template switching in (CAG)_102_ c.13 cells. However, examination of the (CAG)_102_ repeat showed that POLη knockdown increased the breadth of the CAG microsatellite deletion (Figure 14, panels (C) – (E)). Over the entire ES, Polη knockdown increased the breadth of deletions from 6.7 to 8.4 deleted bases/kb, but slightly decreased the number of deletion events from 1.66 to 1.62 deletion events/kb. We conclude that the loss of TLS POLη activity, or depletion of COPS2, destabilizes replication of the (CAG)_102_ microsatellite, but that the gross genomic effects of COPS2 knockdown are not entirely accounted for by POLη depletion. In addition, depletion of COPS2 or POLη changed the base substitution signatures of both (CAG)_102_ clones (Supplementary Figure 8).

**Figure 14:**
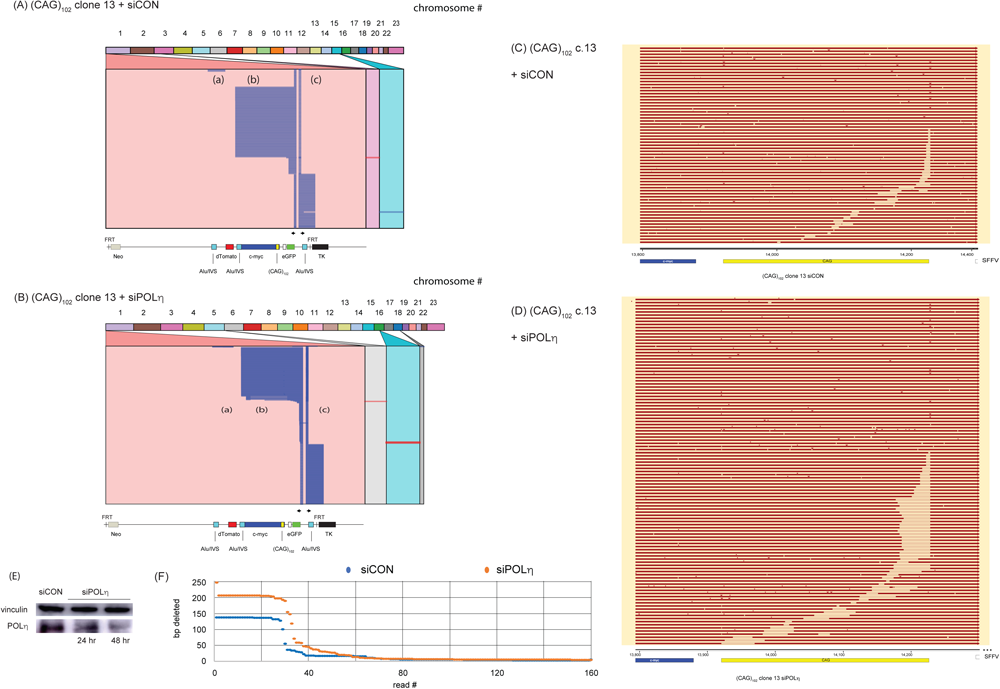
Effect of POLη depletion on eccDNA mutagenesis. Alignment (Ribbon) of iPCR reads from (A) (CAG)102 clone 13 cells, (B) (CAG)102 clone 13 cells depleted of POLη. Alignment (Snapgene) of ectopic site (CAG)102 region in (C) (CAG)102 clone 13 cells, (D) (CAG)102 clone 13 cells depleted of POLη. Alignments are phased based on partial deletions within the CAG repeats. (E) Western blot (CAG)102 clone 13 cells. (F) Depletion of POLη increased the breadth of short deletions from 6.7 to 8.4 deleted bases/kb, p<0.04) at the ectopic site, but slightly decreased the overall number of deletion events at the ectopic site from 1.66 to 1.62 deletion events/kb p<0.028 (not shown)). P values, Student’s t-test

The populations seen after knockdowns or drug treatment began as sister aliquots of a mixed population. Therefore, a change in the appearance of the majority of the population after transient knockdown is not due to a clonally propagated stochastic change.

## Discussion

Extrachromosomal DNAs are found in all species of eucaryotic cells, in a wide range of sizes and compositions. We have used microsatellites capable of forming hairpin, quadruplex, triplex and unwound DNAs to show that unstable microsatellite sequences are centers for the formation of eccDNAs in human chromosomes. The eccDNAs that we have characterized appear to be in the range of ∼250 bp – 15 kb, however, because we used head-to-head primers for iPCR, we do not formally know the length or sequence of the DNA that might be between the apposed 5’ ends of the iPCR primers. Nevertheless, close inspection of several alignments (Figures 4, 11, 13) shows a small number of reads that span the 5’ ends of the primers, which we attribute to rolling circle replication during iPCR. Invariably, these reads contain only the expected ES sequences between the 5’ ends of the primers.

Our data show that the instability of microsatellite non-B DNAs promotes the synthesis of highly mutagenized eccDNAs, and that exposure to replication stress changes the mutagenesis and template switching patterns over the ES eccDNAs. Relevant to ensemble analyses of DNA damage, we show that the same form of stress has different effects on the structure and mutagenic signatures of different microsatellites, and that different forms of replication stress elicit discrete consequences at the same non-B DNA. As well, depletion of the replication factors Rad51, COPS2, or POLη alters the structure and mutagenic signatures of the eccDNAs in distinct ways.

The template switching domains in the eccDNAs are recurrent, but with nonidentical boundaries, and the patterns of overlap microhomology between switching events are strongly nonrandom and microsatellite-specific. We suggest that the avoidance of specific overlap lengths is determined not only by 3’ end microhomology, but also by additional sequences upstream of the 3’ end. This is consistent with suggestions that sequences upstream of the invading 3’ end are involved in the mechanism of selecting among multiple potential invasion sites (115,117). We also speculate that the same DNA damage tolerance polymerase may be responsible for successive template switch overlaps in individual reads.

Autonomous replication activity has been attributed to double minute chromosomes (152) and to random DNA sequences dependent on their length (98). We expect that a fraction of the eccDNAs that we have identified may have this ability, and note that the (ATTCT)_47_ and (CAG)_102_ c.10 cell lines display distinct populations of yellow cells with enhanced fluorescence, consistent with the presence of extrachromosomal copies of the dTomato gene or the eGFP gene. The putative ability of eccDNAs to replicate autonomously supports the possibility of ongoing replication-dependent mutation in these molecules.

Expanded CTG/CAG repeats form hairpins, stall replication forks, and are targets for DSBs in vivo (59,67). In agreement with a model of BIR from a deDSB, mutagenesis occurs upstream and downstream from DSBs at CTG/CAG microsatellites in the presence of replication stress, dependent on POLD3 and BRCA2 (22). In addition, mutagenesis and genome instability occur upstream and downstream of the Pu/Py mirror repeat polar replication barrier which recruits the DNA damage response proteins ataxia telangiectasia mutated and Rad3-related (ATR) and Rad9 (77).

H3 and G4 sequences derived from the Pu/Py repeat also elicit gross chromosomal instability, and ligand-induced G4 non-B DNA induces mutagenesis at the upstream dTom and downstream eGFP reporters (21). Similarly, expanded ATTCT repeats induce hypermutagenesis and chromosome instability (30). Consistent with evidence that knockdown of the signalosome component COPS2 (20), Rad18, or the TLS polymerases POLη and POLκ, increase microsatellite instability (20,22), we show here that replication stress due to HU, APH or depletion of COPS2 , POLη or Rad51 markedly affect eccDNA abundance and structure.

The prolonged lifetime of single stranded DNA during lagging strand replication or BIR is believed to favor the formation of non-B DNA structures, including G quadruplexes (42,80,153,154). In human cells G quadruplexes form preferentially on the lagging strand replication template, and induce DNA strand breaks and BIR (21,22). In yeast, however, there are conflicting reports of the template preference for quadruplex formation under specific conditions (155,156). Consistent with the slow rate of BIR and the sustained ssDNA of the D-loop and nascent BIR DNA, the results of Figures 4 implicate G quadruplex forming sequences as frequent sites of instability during BIR and eccDNA formation.

Within the same parent DNA sequence (e.g. nonmicrosatellite segments of the ES), the pattern of template switching and the locations of mutations differ significantly between reads within a clone, and between sister or nonsister clones. Taking into account the number of population doublings in each of our clonal cell lines, we calculate mutagenesis rates in the eccDNAs (indel events plus substitutions) of ∼3.4 x 10^-6^/generation – 1.4 x 10^-5^/generation. These mutations are found both upstream and downstream of the ectopic microsatellite sequence and the ectopic c-myc origin. The pattern and abundance of mutations suggest a bidirectional mechanism of BIR.

In *S. cerevisiae*, DSBs are repaired by long tract gene conversion (LTGC) and synthesis-dependent strand annealing (SDSA) if both ends have proximal homology on the sister chromatid (54,55,157), whereas mutagenic BIR from a single-ended DSB (seDSB) can occur if a parental strand at the DSB becomes ligated to a nascent DNA strand. Alternatively, bidirectional BIR can occur if the ends do not contain proximal (∼1-2 kb) homology on the donor DNA (131,157,158), or when the second end of the break is not captured (131,159,160) due to a converging fork, the absence of proteins that promote strand annealing (161) or due to singular structures of DNA or chromatin (103,162–173).

We have shown that replication of the ES initiates from the proximal c-myc origin in >90% of S-phases, that discrete DSBs occur at the downstream edge of the expanded (CTG/CAG) microsatellite (22), and that mutagenesis occurs upstream and downstream of a G4 forming sequence at the same ES (21). In the present work, sequence analysis reveals template switches, indels and substitution mutations extending >5-10 kb upstream and downstream of the ectopic ATTCT, CAG, H3 and G4 microsatellites within individual sequence reads. In addition, flow cytometry shows cells that delete or mutagenize both reporter genes, or exclusively lose expression of the upstream dTomato marker or the downstream eGFP marker. Taken together, these observations are consistent with a role for BIR from a deDSB between the reporter genes in the generation of highly mutagenized eccDNAs.

There are certain limitations to our study. First, the maximum size of our iPCR products is ∼15 kb, and due to the iPCR strategy, we do not have information on DNA between the 5’ termini of the inverse primers. Therefore unknown DNA sequences could be present in the eccDNAs. We note, however, that in those cases where the Q5 polymerase has apparently copied the template by a rolling circle mechanism, the sequences between the primer 5’ ends are the expected ES sequences. Second, our data cannot be used to quantitate eccDNAs, due to the variability of PCR efficiency, and because of the unpredictability of eccDNA structures.

Based on our data we propose an ‘asynchronous capture’ mechanism as an additional choice for the repair of replication-dependent microsatellite DNA damage (174,175), in which endonuclease cleavage at a stalled replication fork results in bidirectional BIR (Figure 15). The model predicts that replication through the microsatellite, and from a converging origin, produce a deDSB (panel (A)). Delayed 5’ resection of one end (panel (B)) leads to asynchronous capture of the 3’ ends by the sister chromatid or nonallelic chromosomes, and bidirectional BIR (panel (C)). Anti-recombinase/helicase displacement of the invading strands allows the completion of conservative DNA synthesis (panel (D), resulting in two chromosome fragments, each carrying a telomere end and a ‘BIR end’ (panel (D)). Homologous recombination (panel (E)) yields two crossover structures, one of which is a chromosome containing two telomeres and a homologous recombination deficiency (HRD scar (119,135,136,176), and the other which contains two ‘BIR ends’. Homologous recombination at internal repeated ‘eccDNA junction’ sequences produce circular DNAs. Experiments are underway to test this hypothesis.

**Figure 15.**
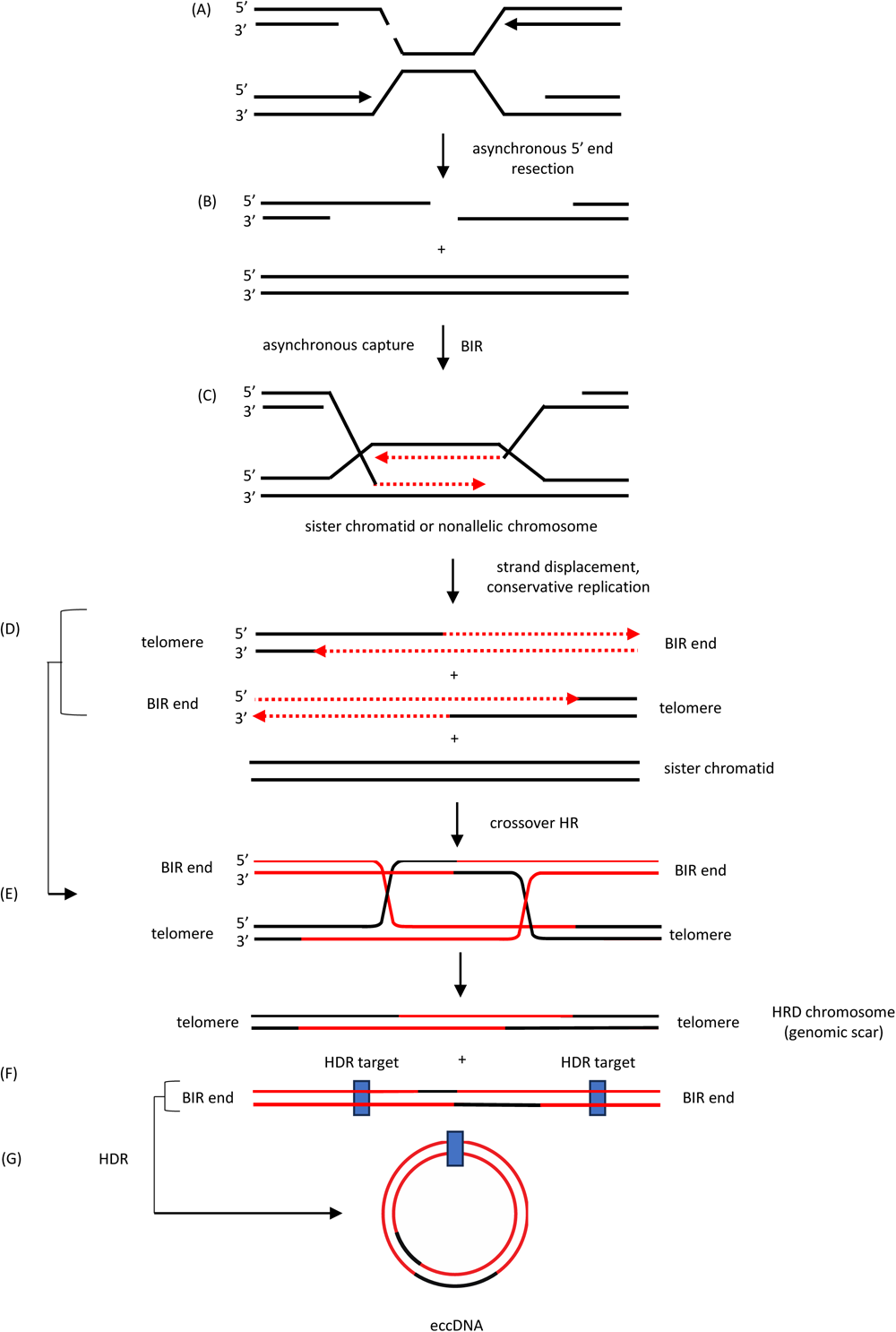
Proposed asynchronous capture model of BIR. (A) In this model the rightward replication fork initiates at the c-myc origin (22), and the leftward fork initiates 8-20 kb downstream (58) to converge at the double strand break at non-B microsatellite DNA. (B) Asynchronous resection or capture of the DSB ends leads to (C) bidirectional BIR and template switching on the sister chromatid, the self chromosome (not shown) or a nonallelic chromosome. (D) Anti-recombinase/helicase activity allows displacement of the nascent DNA strands and completion of conservative replication, restoring the sister chromatid and producing two incomplete chromosomes, each with one telomere and one end determined by the extent of BIR (“BIR end”). (E) Crossover homologous recombination results in one chromosome with two telomeric ends and an internal HRD ‘genomic scar’ and a second chromosome with two BIR ends. (F), (G) The chromosome fragment with two BIR ends undergoes homology-dependent recombination at one or more repeated sequences to produce eccDNA(s).

## Data Availability

PacBio sequence data and HeLa/GRCh38/ectopic site reference genome data are available at the NCBI Sequence Read Archive (SRA) link: https://www.ncbi.nlm.nih.gov/bioproject/PRJNA971563. Additional data are available upon request from the authors.

## Acknowledgments

We thank Michael Markey (WSU Center for Genomics Research) for assistance with flow cytometry and Michael Markey, Yong-Jie Xu and Michael Kemp for comments on this work, and Jin Mo Park (Massachusetts General Hospital) for suggesting a role for recombined eccDNAs in neoantigen synthesis.

## Support

This work was supported by the NIH [grant GM122976 to ML]. VA and RS also thank the Wright State University Biomedical Sciences Ph.D. Program for support. The content of this work is solely the responsibility of the authors and does not necessarily represent the official views of the National Institutes of Health.

## Conflict of Interest Statement

The authors declare no conflicts of interests related to this work.

## Notes

### Competing Interest Statement

The authors have declared no competing interest.

