## Supplementary Figure 1 for "Microsatellite break-induced replication generates highly mutagenized extrachromosomal circular DNAs"

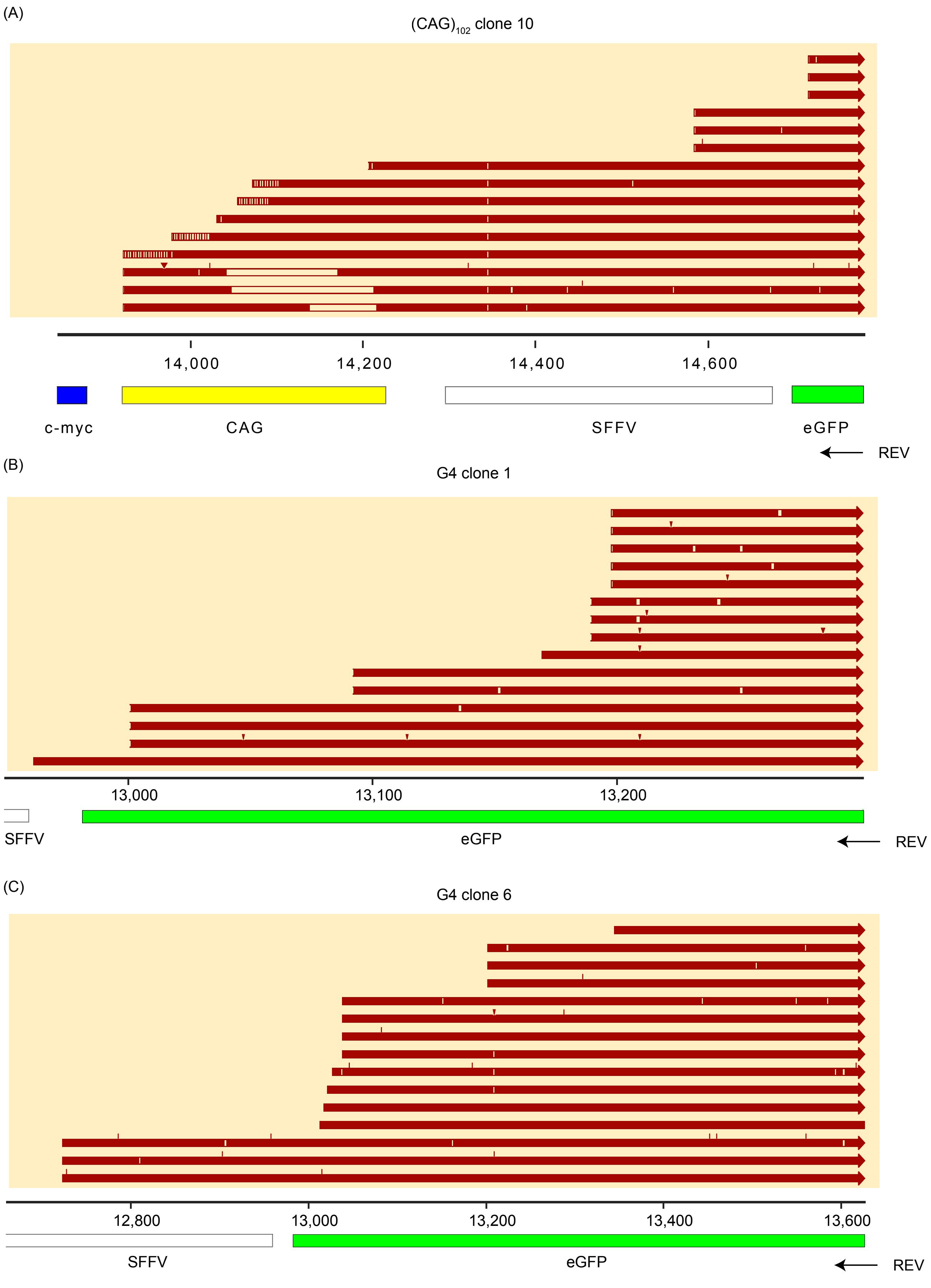
(A)
(CAG)102 clone 10
14,000
14,200
14,400
14,600
c-myc
CAG
SFFV
eGFP
REV
(B)
G4 clone 1
13,000
13,100
13,200
SFFV
eGFP
REV
(C)
G4 clone 6
12,800
13,000
13,200
13,400
13,600
SFFV
eGFP
REV

**Supplementary Figure 1. Variable boundaries of template switching domains. The 3' ends of the reverse iPCR primer reads were mapped against segments of the ectopic site in (A) (CAG)102 c.10 cells, (B) G4 c.1 cells, (C) G4 c.6 cells. Open boxes represent deletions; vertical bars (and inverted triangles) represent single base (or multiple base) mismatches and insertions.**
