## Supplementary Figure 2 for "Microsatellite break-induced replication generates highly mutagenized extrachromosomal circular DNAs"

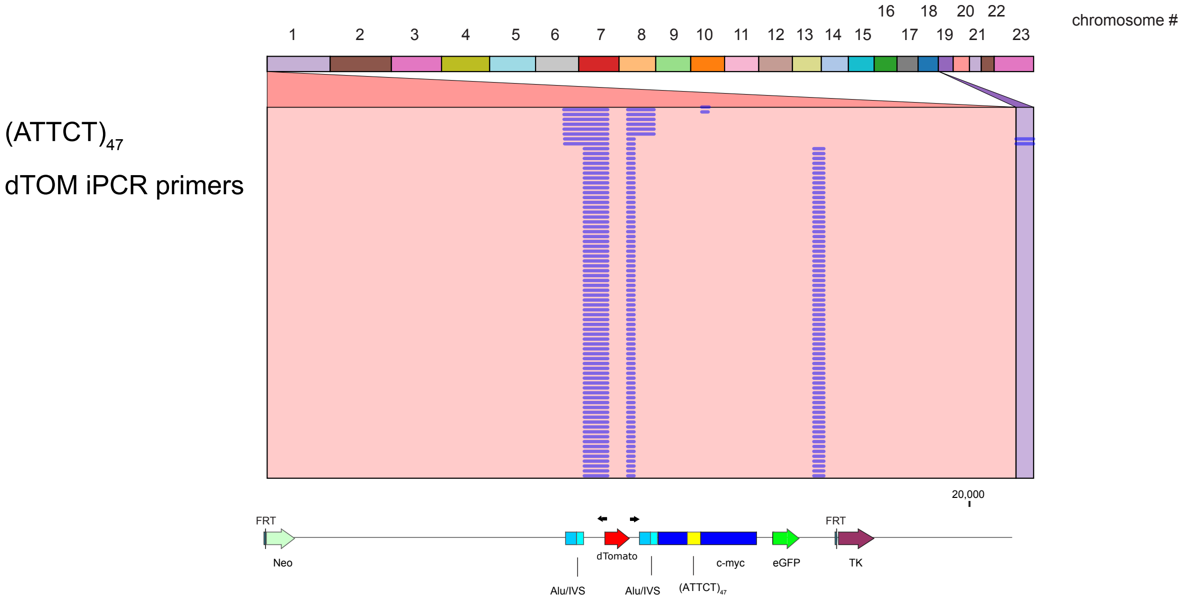


**Supplementary Figure 2. Multiple eccDNAs derive from a single ectopic site. iPCR primers flanking the dTOM gene were used to generate 220 reads from (ATTCT)47 cells, which are aligned to the ectopic site and chromosome 19. The template switch to chromosome 19 (nt 2,010,376 – 2,010,938) occurred in a 562 bp region with four G4 consensus matches.**
