## Supplementary Figure 3 for "Microsatellite break-induced replication generates highly mutagenized extrachromosomal circular DNAs"

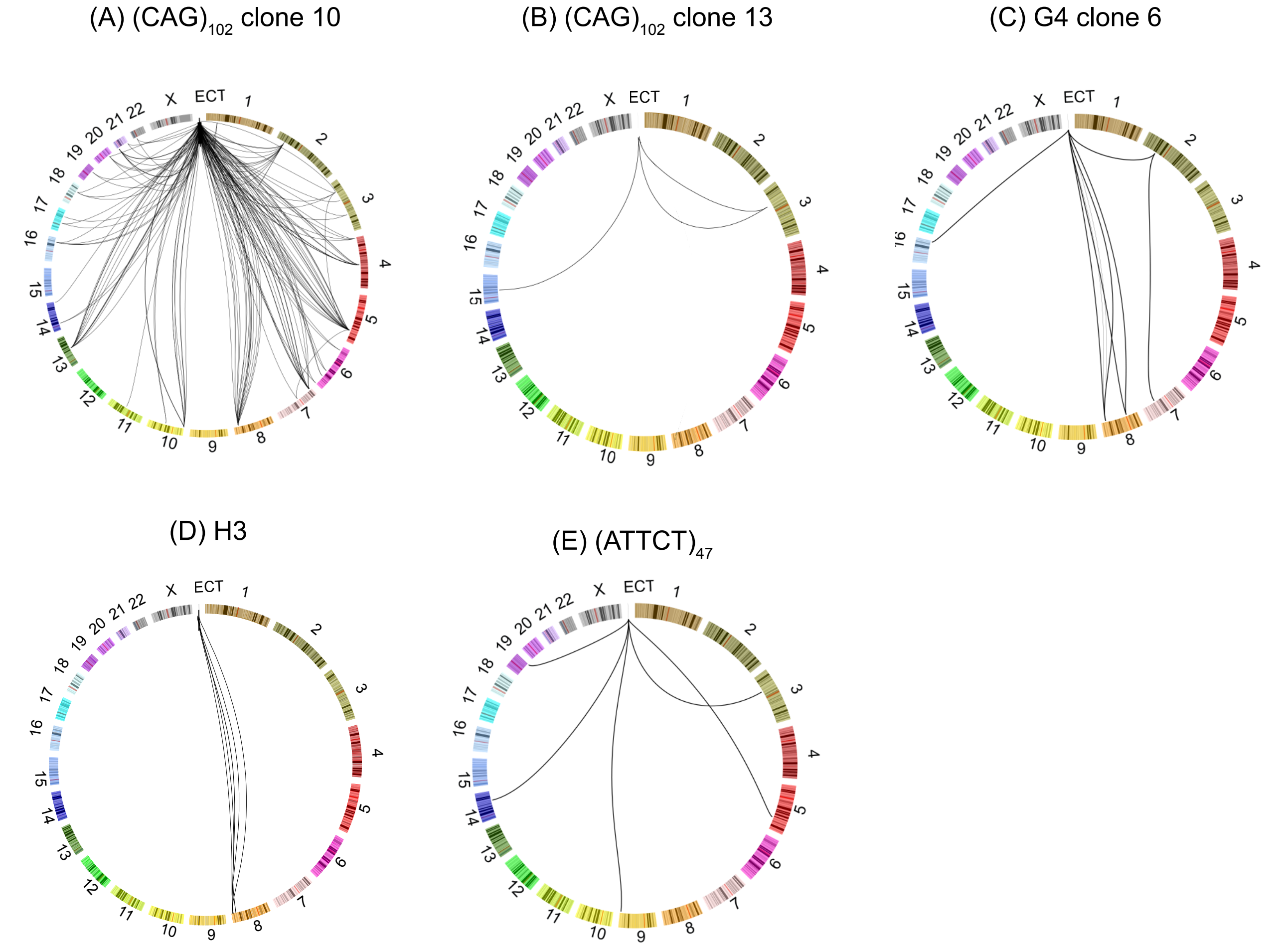


**Supplementary Figure 3. Nonallelic template switches. Nonallelic template switches in eccDNAs were visualized using Circos (64). (A) (CAG)102 c.10 cell eccDNA, (B) (CAG)102 c.13 cell eccDNA, (C) G4, c.6 cell eccDNA, (D) H3 cell eccDNA, (E) (ATTCT)47 cell eccDNA; ECT, ES.**
