## Supplementary Figure 4 for "Microsatellite break-induced replication generates highly mutagenized extrachromosomal circular DNAs"

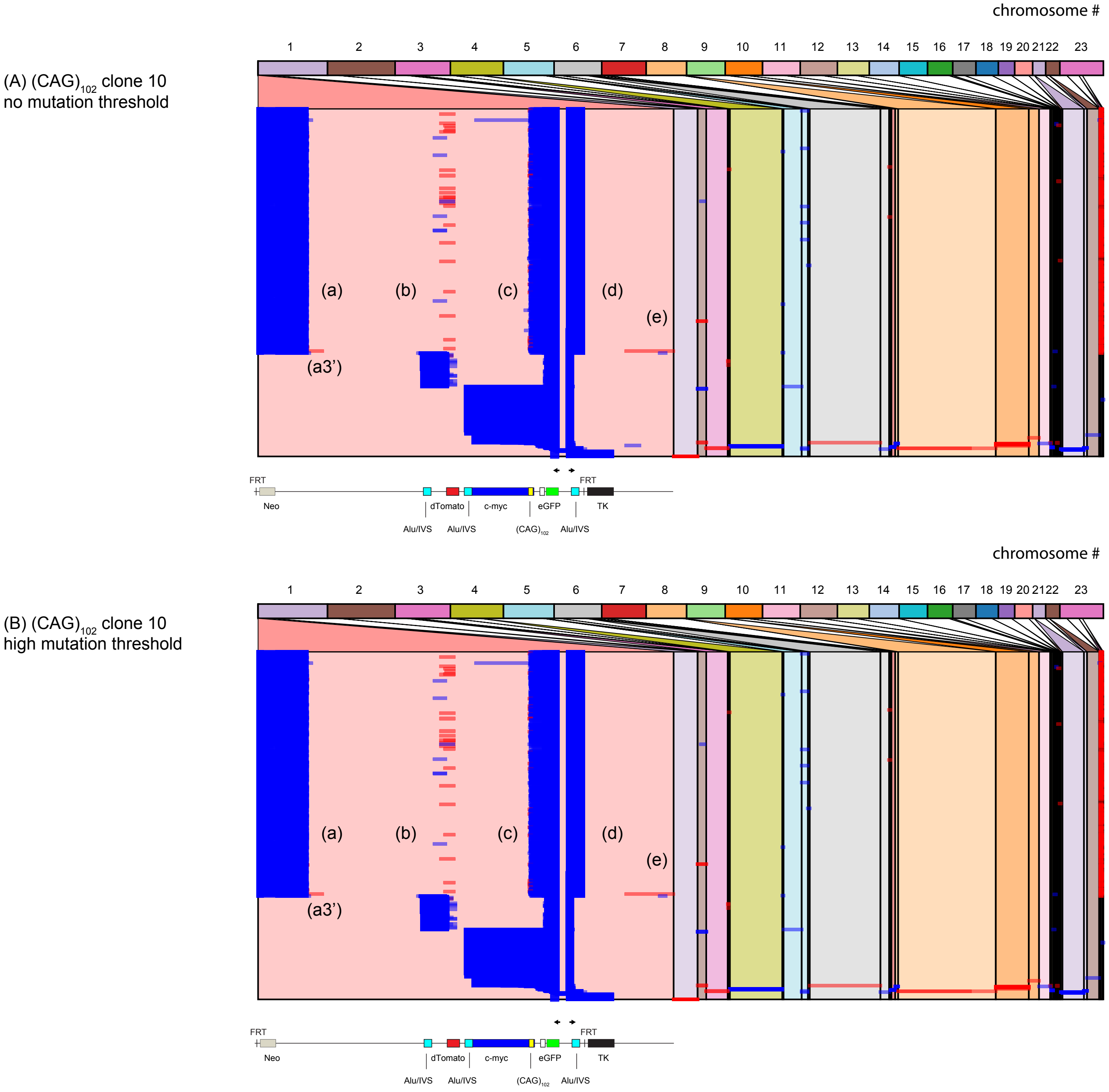

Supplementary Figure 4

(C) $(CAG)_{102}$ clone 13
no mutation threshold

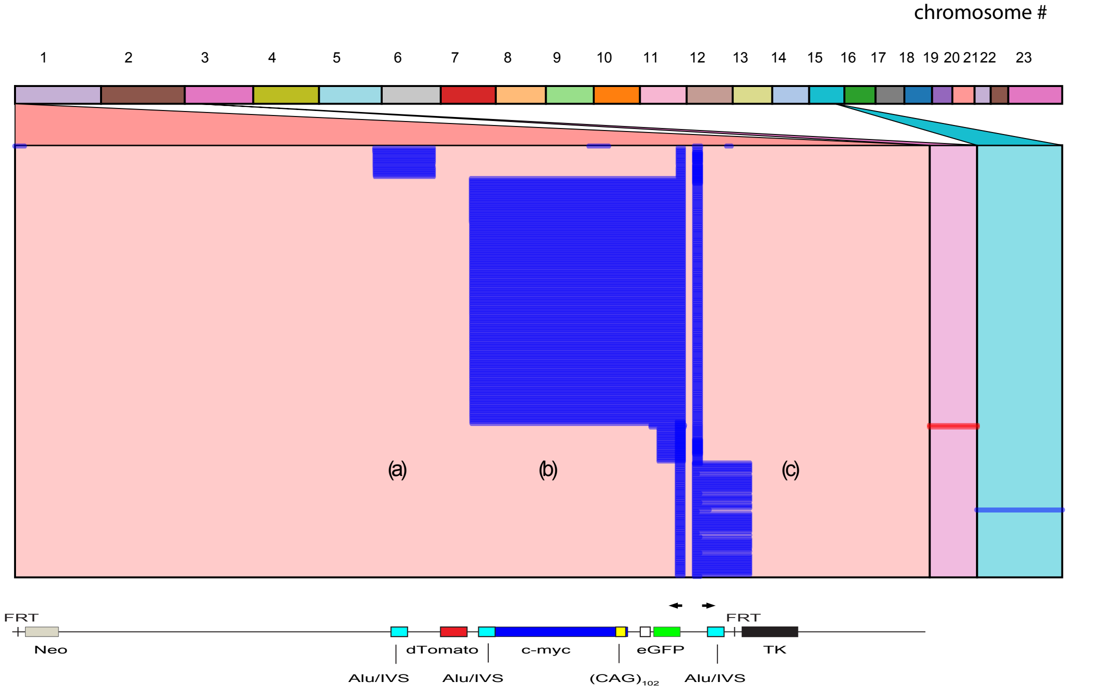


(D) $(CAG)_{102}$ clone 13
hgh mutation threshold

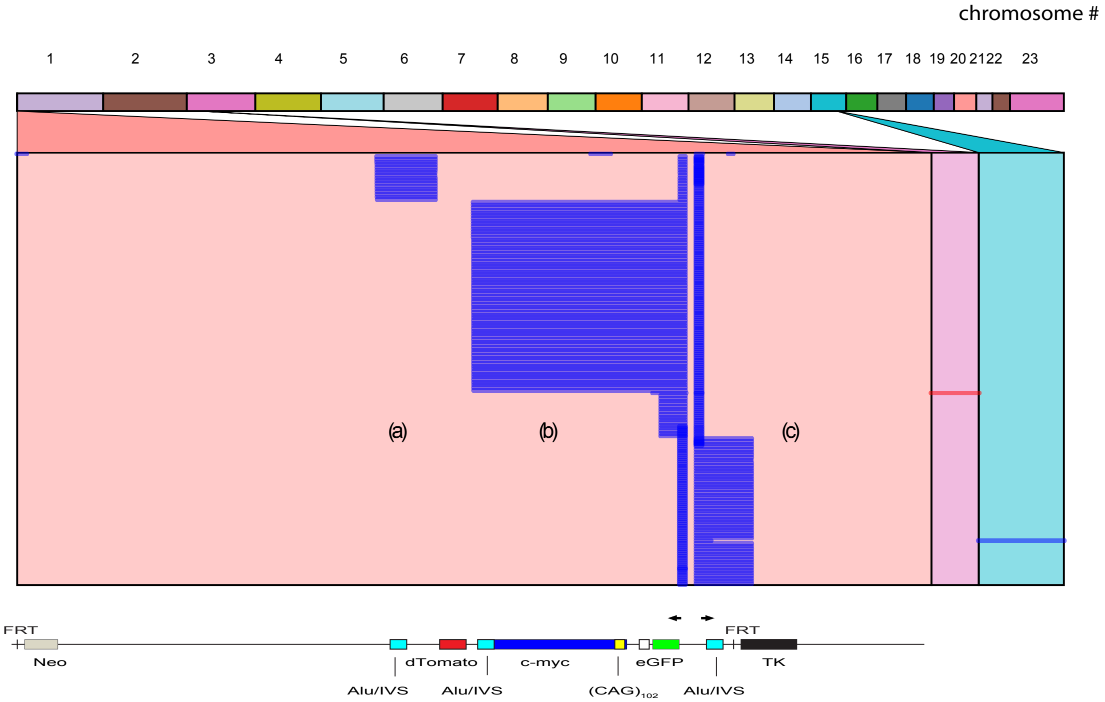

**Supplementary Figure 4. Application of a high mutation threshold to (CAG)102 eccDNA alignments. Alignment of (CAG)102 clone 13 reads without (C) or with (D) a mutation of threshold of 2.0 mutations (indels, base substitutions) per kb.**
