## Supplementary Figure 5 for "Microsatellite break-induced replication generates highly mutagenized extrachromosomal circular DNAs"

Figure 5A

(CAG)$_{102}$ clone 13

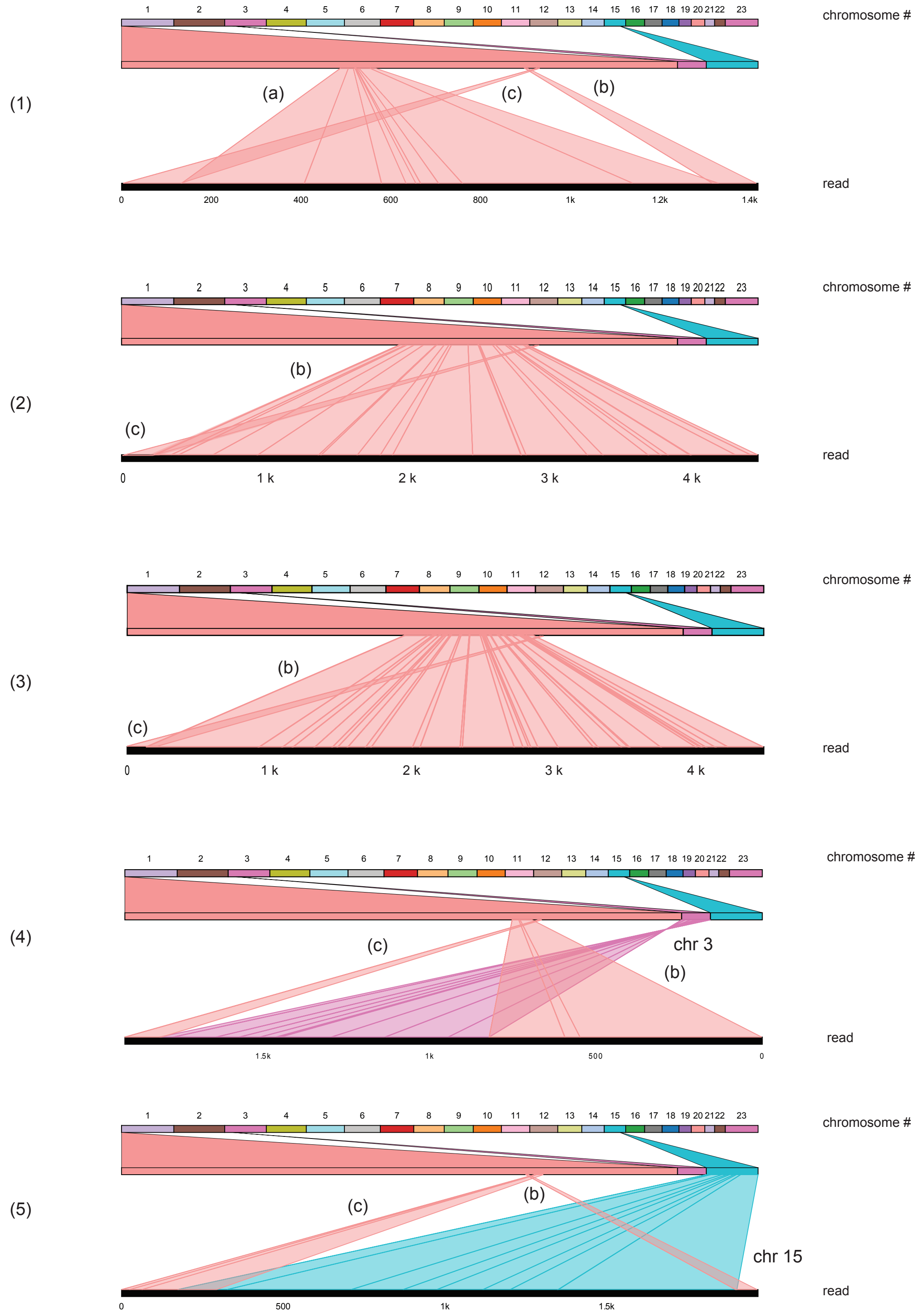

Supplementary Figure 5B

### G4 clone 1

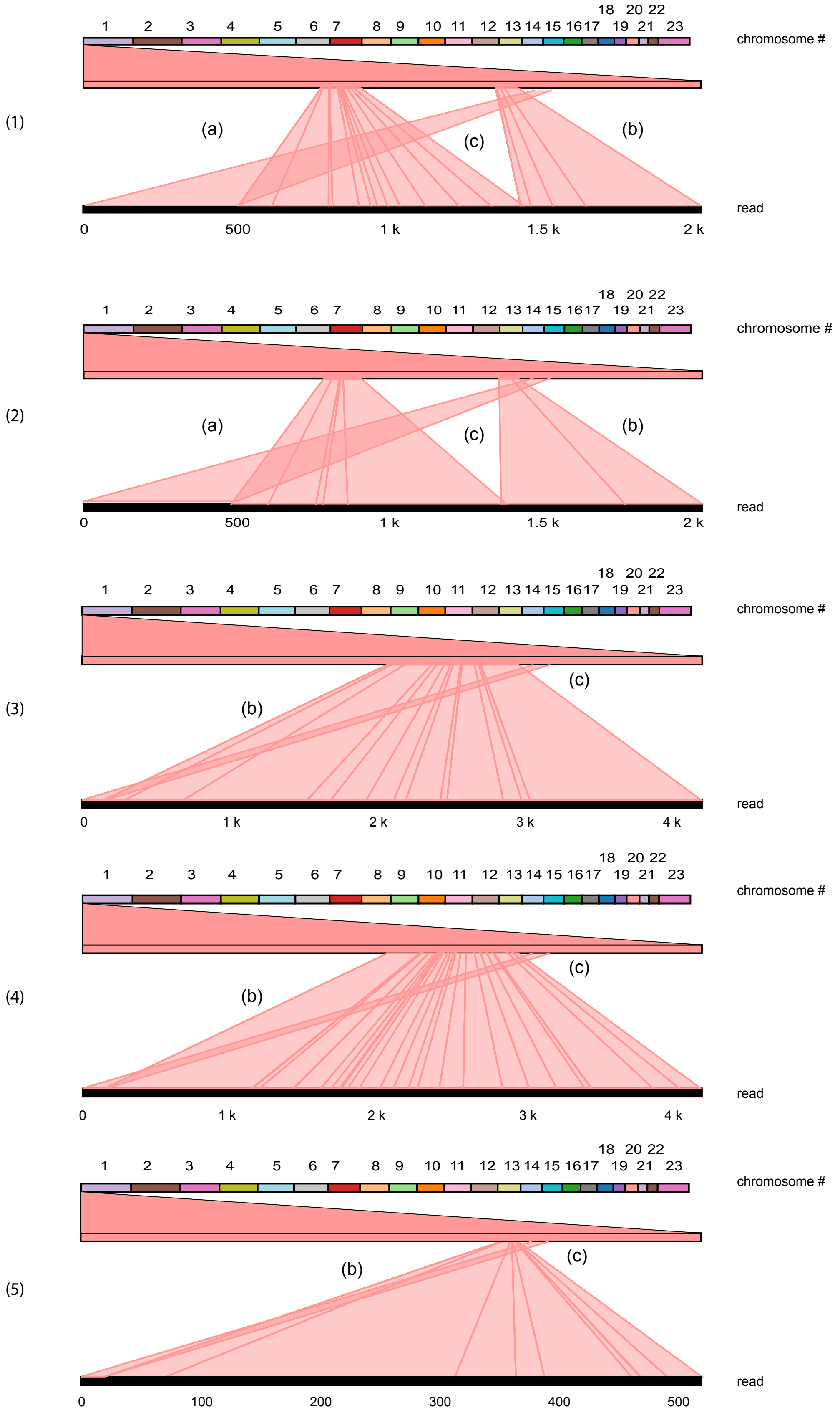

G4 clone 6

Supplementary Figure 5C

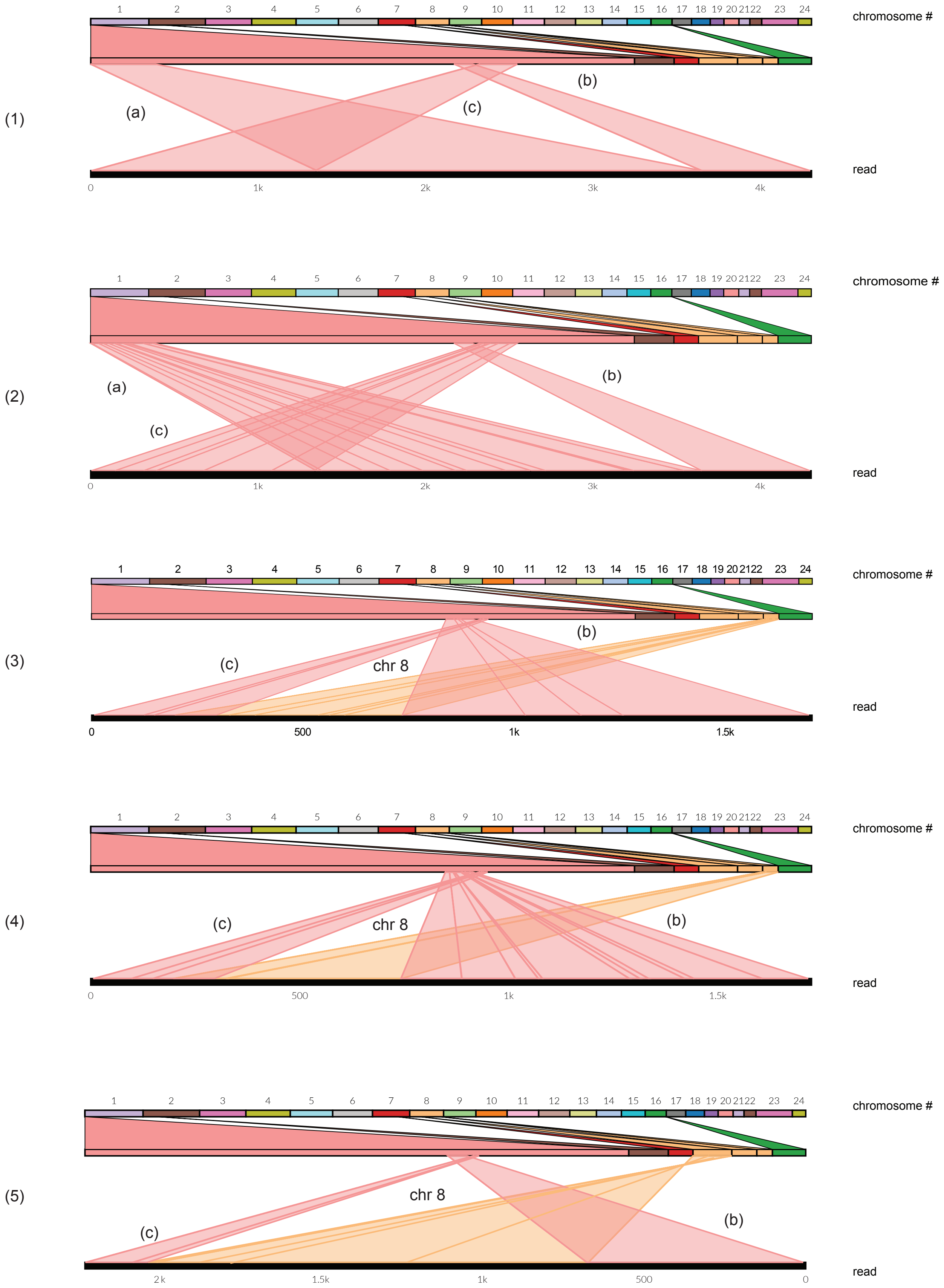

Supplementary Figure 5D

## H3

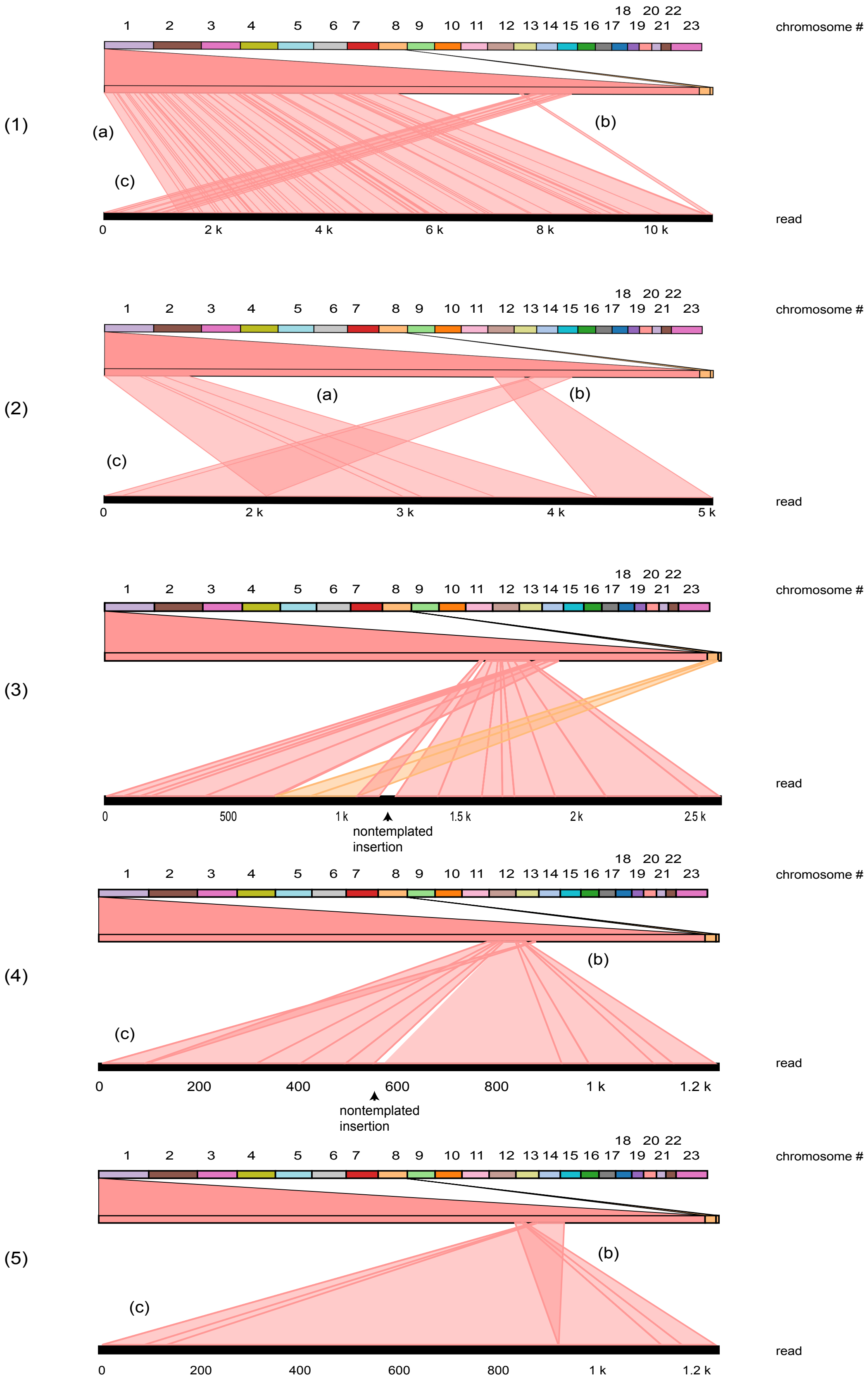

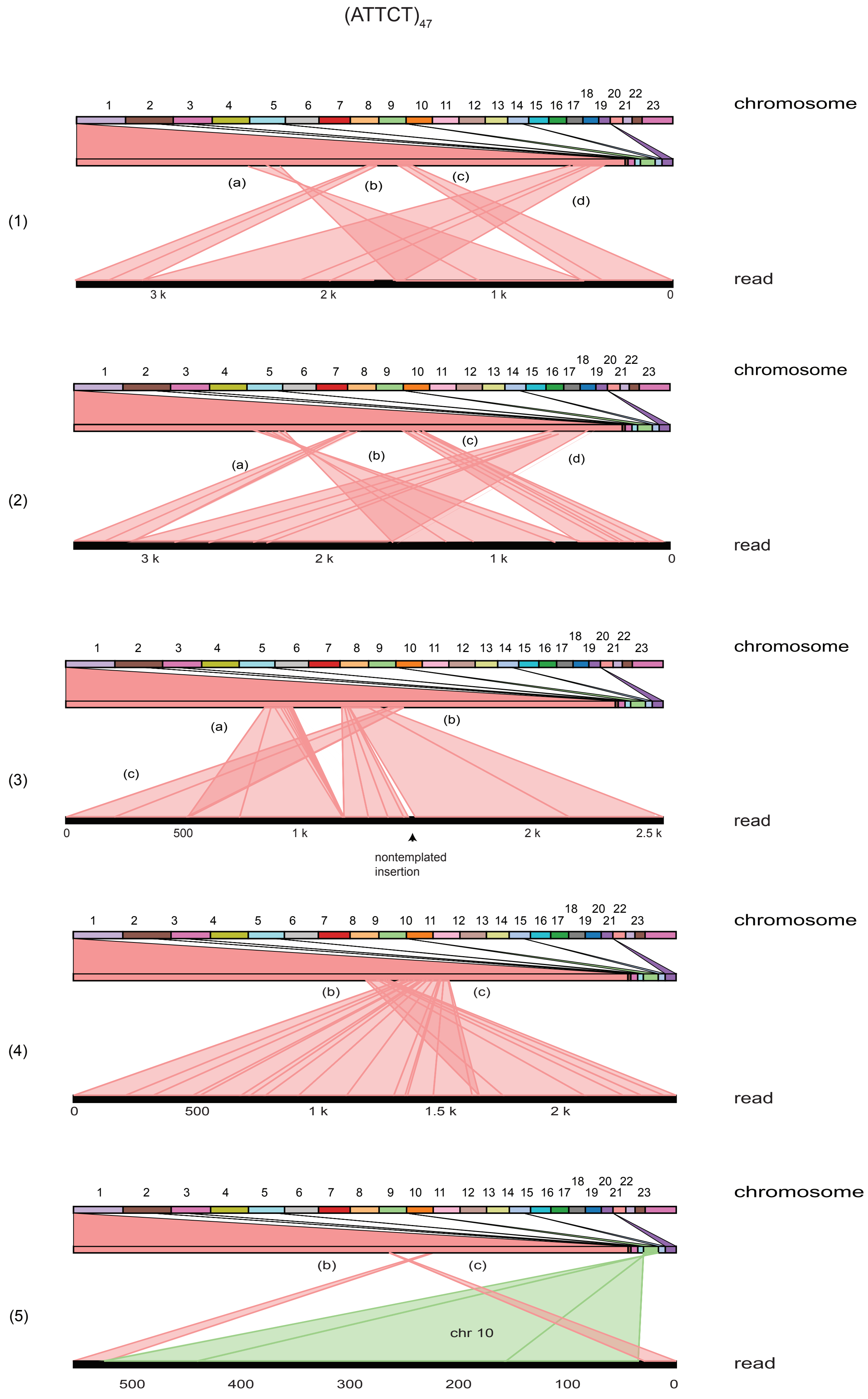
(ATTCT)$_{47}$
chromosome
read
(1)
(a)
(b)
(c)
(d)
3 k
2 k
1 k
0
(2)
(3)
500
2.5 k
nontemplated insertion
(4)
1.5 k
(5)
chr 10
400
300
200
100

**Supplementary Figure 5: Template switching patterns of eccDNAs in (CAG)102 clone 13. (A) Individual reads from (CAG)102 c.13 cells (lines (1)-(5), Figure 4B) were mapped in the query viewport view of Ribbon (61). Chromosome numbers are shown at the top; the lower heavy black line in each panel is the complete read. Heavier red lines within a domain indicate indels. Letters (a), (b), (c), (d) correspond to template switching domains in Figure 4B. (B) Reads from G4 c.1 cells (lines (1)-(5) Figure 4(C)). (C) Reads from G4 c.6 cells (lines (1)-(5) Figure 4(D)). (D) Reads from H3 cells (lines (1)-(5) Figure 4(E)). (E) Reads from (ATTCT)47 cells (lines (1)-(5) Figure 4(F)).**
