## Supplementary Figure 6 for "Microsatellite break-induced replication generates highly mutagenized extrachromosomal circular DNAs"

Supplementary Figure 6A

(CAG)$_{102}$ clone 13

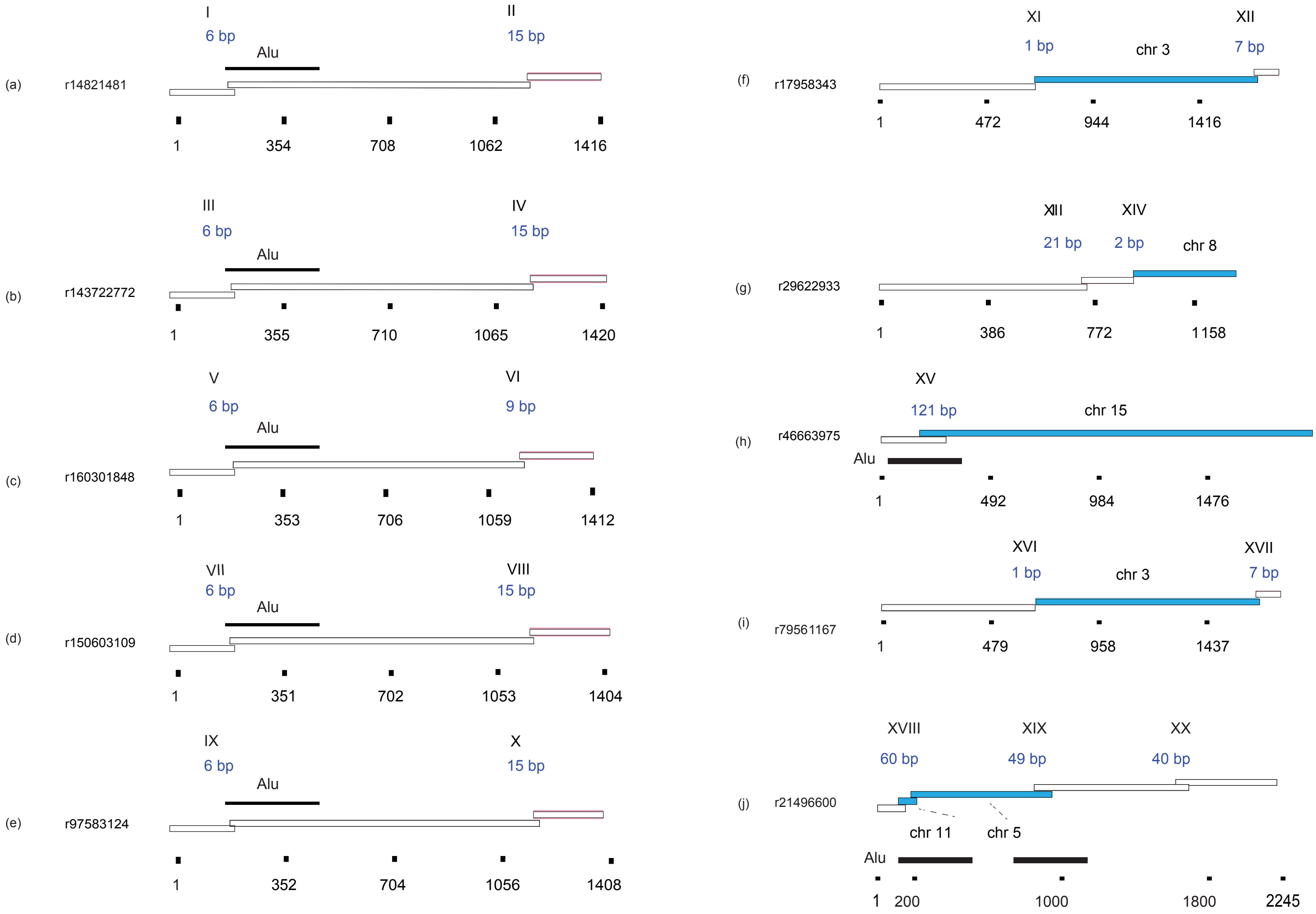

Supplementary Figure 6B

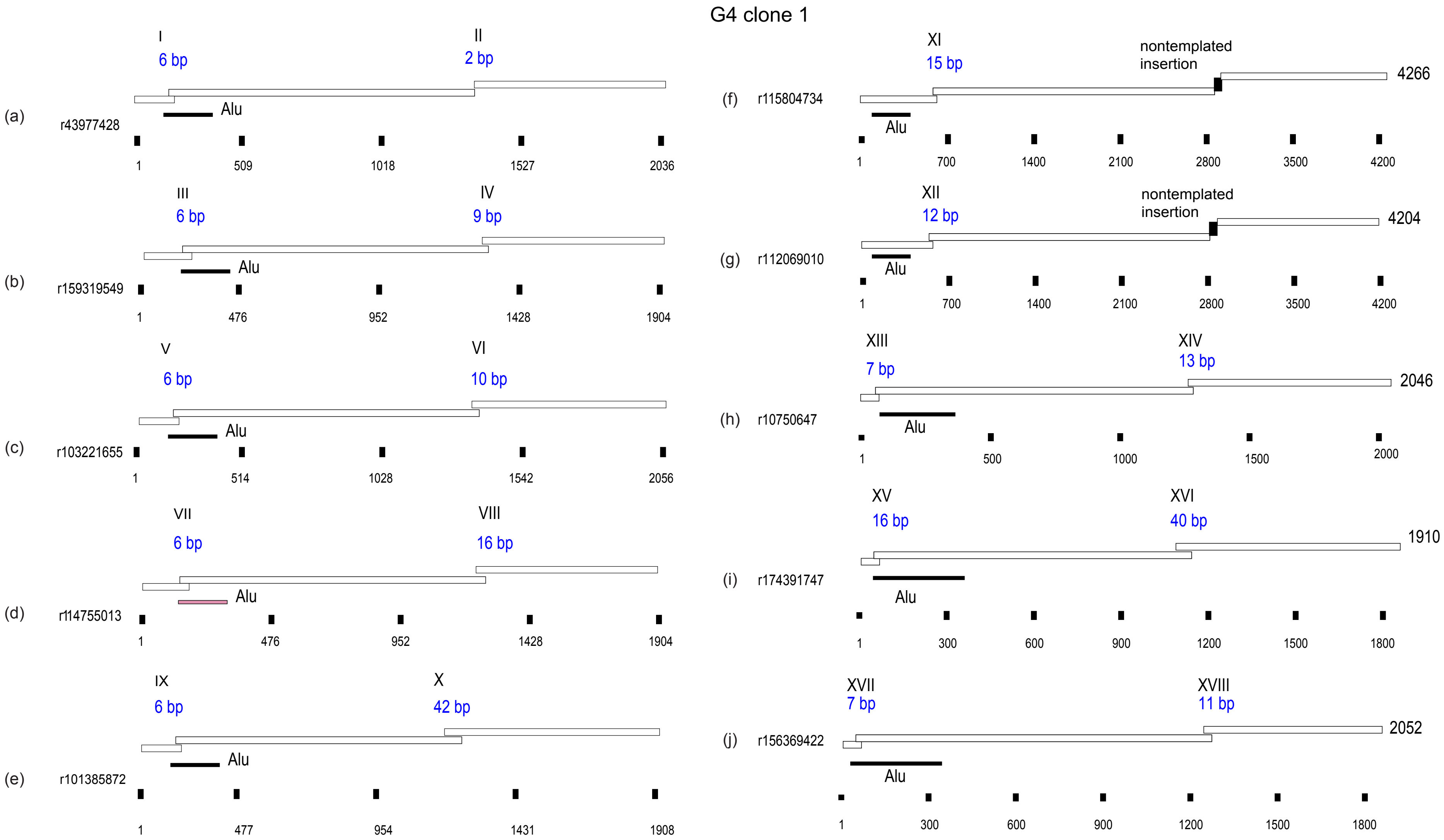

G4 clone 6

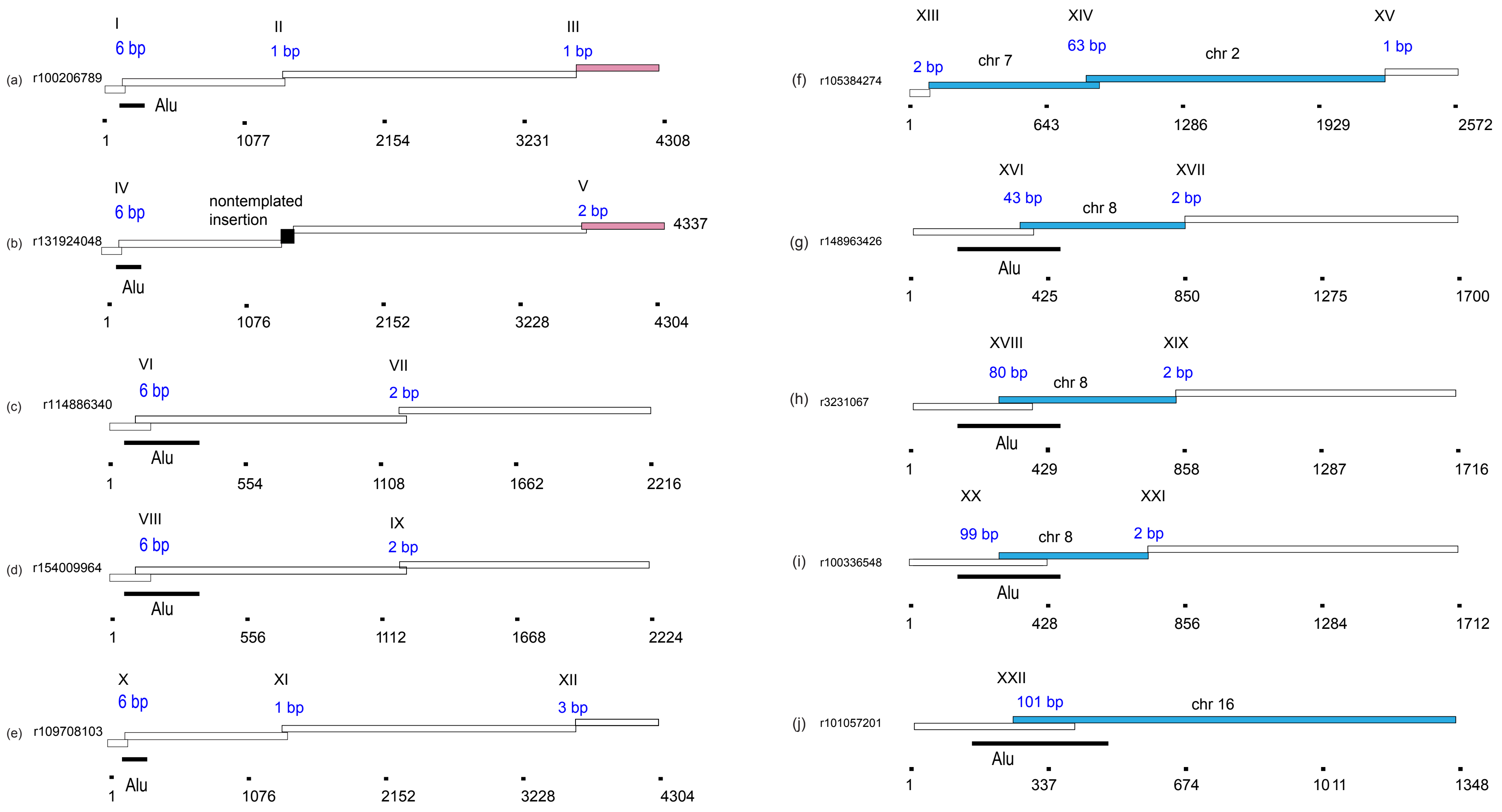

(a) r100206789
I
6 bp
II
1 bp
III
1 bp
Alu
1
1077
2154
3231
4308
(b) r131924048
IV
6 bp
nontemplated insertion
V
2 bp
4337
Alu
1
1076
2152
3228
4304
(c) r114886340
VI
6 bp
VII
2 bp
Alu
1
554
1108
1662
2216
(d) r154009964
VIII
6 bp
IX
2 bp
Alu
1
556
1112
1668
2224
(e) r109708103
X
6 bp
XI
1 bp
XII
3 bp
Alu
1
1076
2152
3228
4304
(f) r105384274
XIII
2 bp
chr 7
XIV
63 bp
chr 2
XV
1 bp
1
643
1286
1929
2572
(g) r148963426
XVI
43 bp
chr 8
XVII
2 bp
Alu
1
425
850
1275
1700
(h) r3231067
XVIII
80 bp
chr 8
XIX
2 bp
Alu
1
429
858
1287
1716
(i) r100336548
XX
99 bp
chr 8
XXI
2 bp
Alu
1
428
856
1284
1712
(j) r101057201
XXII
101 bp
chr 16
Alu
1
337
674
1011
1348

Supplementary Figure 6D

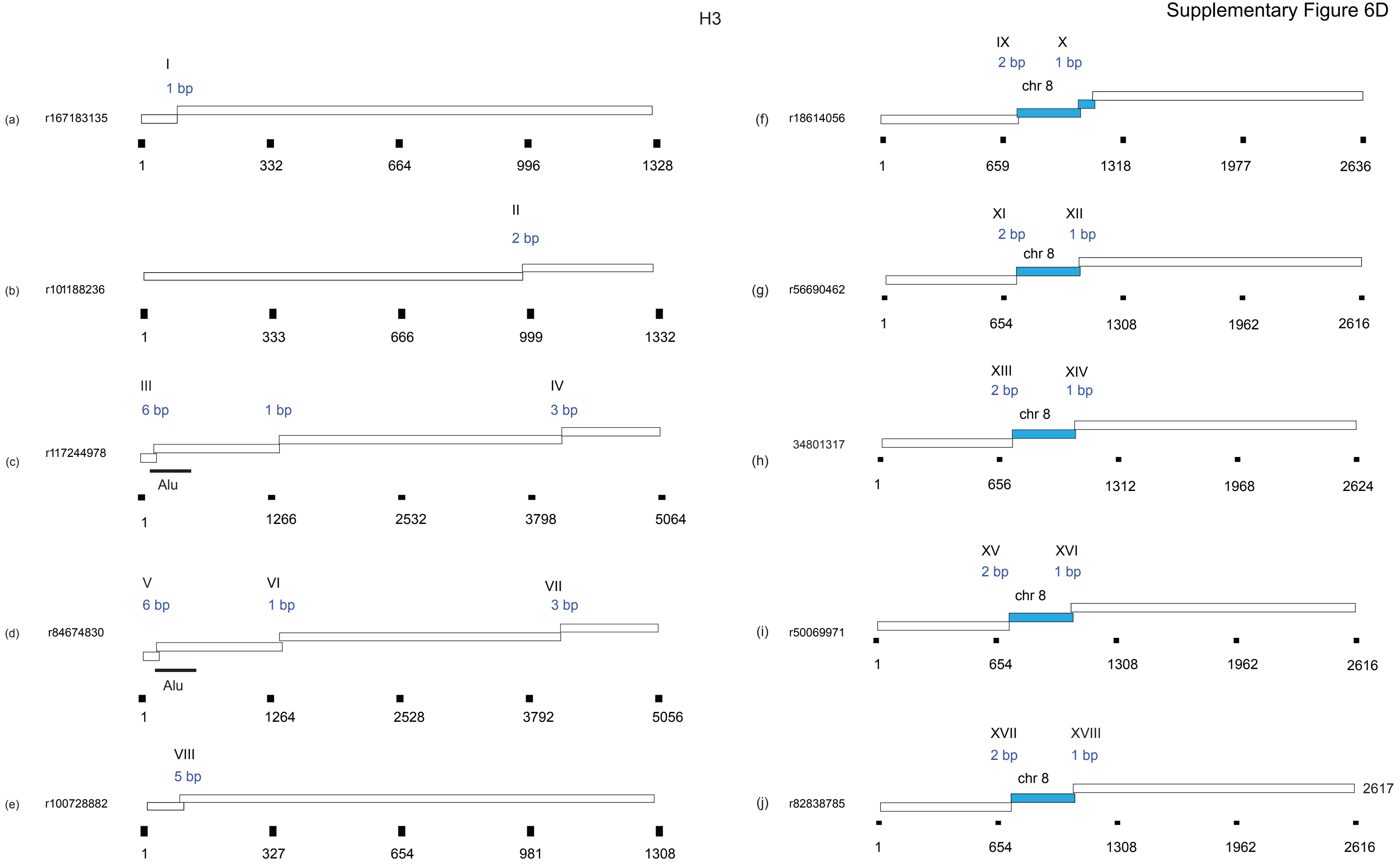

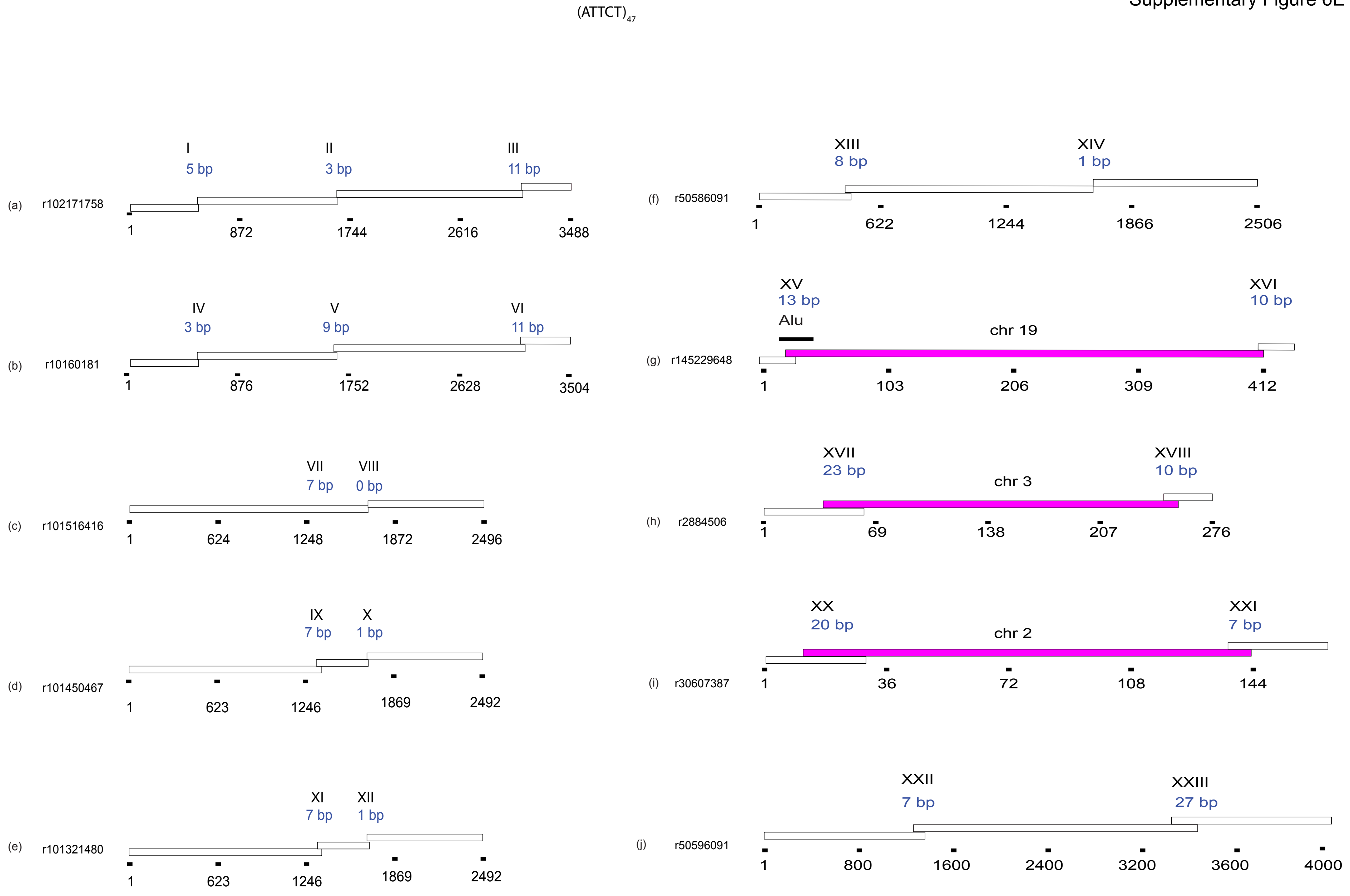
(ATTCT)47
(a) r102171758
I 5 bp
II 3 bp
III 11 bp
1 872 1744 2616 3488
(b) r10160181
IV 3 bp
V 9 bp
VI 11 bp
1 876 1752 2628 3504
(c) r101516416
VII 7 bp
VIII 0 bp
1 624 1248 1872 2496
(d) r101450467
IX 7 bp
X 1 bp
1 623 1246 1869 2492
(e) r101321480
XI 7 bp
XII 1 bp
1 623 1246 1869 2492
(f) r50586091
XIII 8 bp
XIV 1 bp
1 622 1244 1866 2506
(g) r145229648
XV 13 bp
Alu
chr 19
XVI 10 bp
1 103 206 309 412
(h) r2884506
XVII 23 bp
chr 3
XVIII 10 bp
1 69 138 207 276
(i) r30607387
XX 20 bp
chr 2
XXI 7 bp
1 36 72 108 144
(j) r50596091
XXII 7 bp
XXIII 27 bp
1 800 1600 2400 3200 3600 4000

**Supplementary Figure 6: Template switching microhomology. (A) (a)-(j) are individual reads from (CAG)102 clone 13 cells. (B) Reads from G4 clone 1 cells. (C) Reads from G4 clone 6 cells. (D) Reads from H3 cells. (E) Reads from (ATTCT)47 cells. Each box is a template switch domain. Blue domains are nonallelic template switches. Roman and blue bp numerals are sequence overlaps. Approximate length coordinates are shown below or alongside each read. Analysis was performed with ALVIS (178), and reformatted for figure presentation.**
