## Supplementary Figure 7 for "Microsatellite break-induced replication generates highly mutagenized extrachromosomal circular DNAs"

Supplementary Figure 7A

(CAG)$_{102}$ clone 10 plus HU

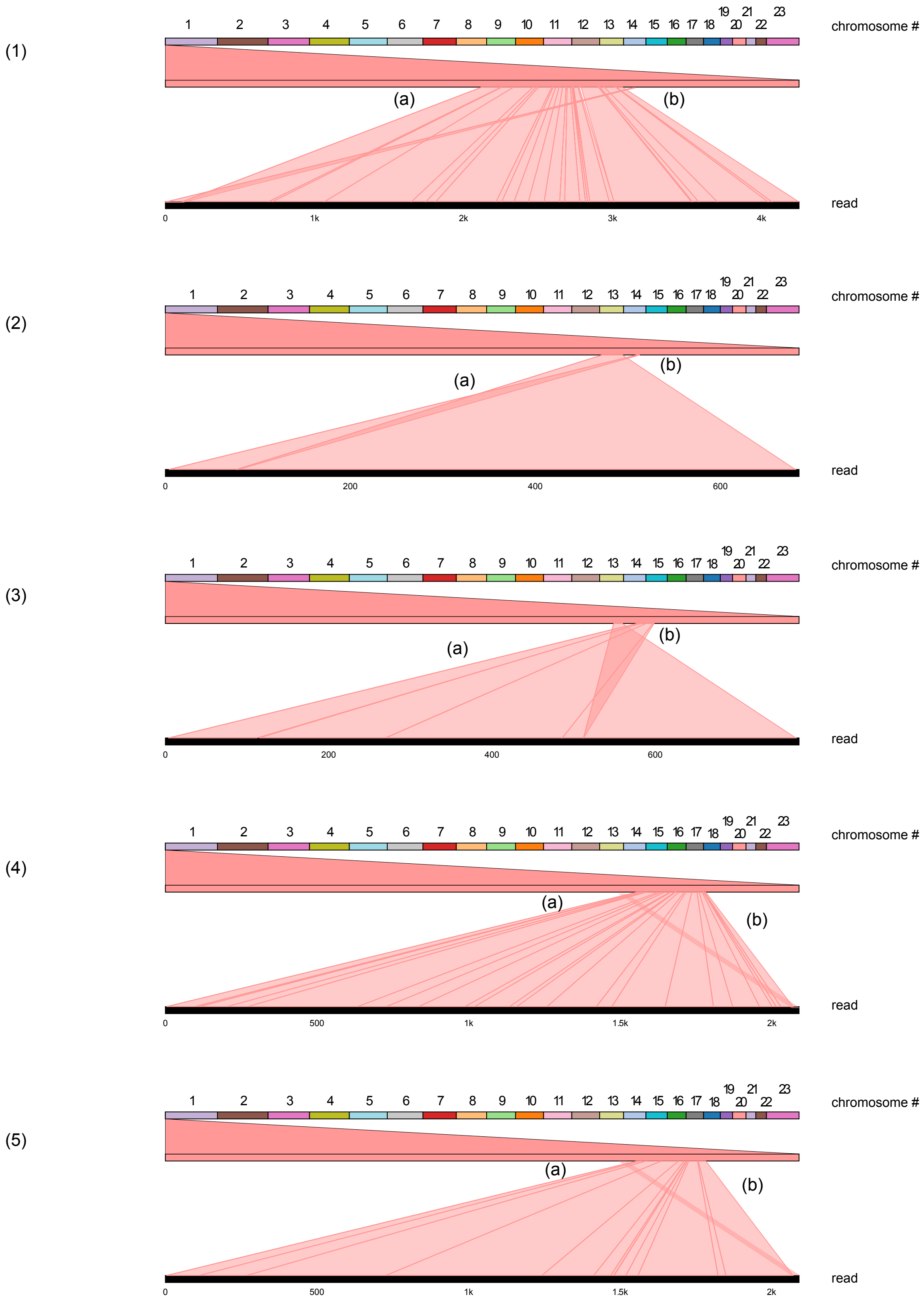

### $(CAG)_{102}$ clone 10 plus APH

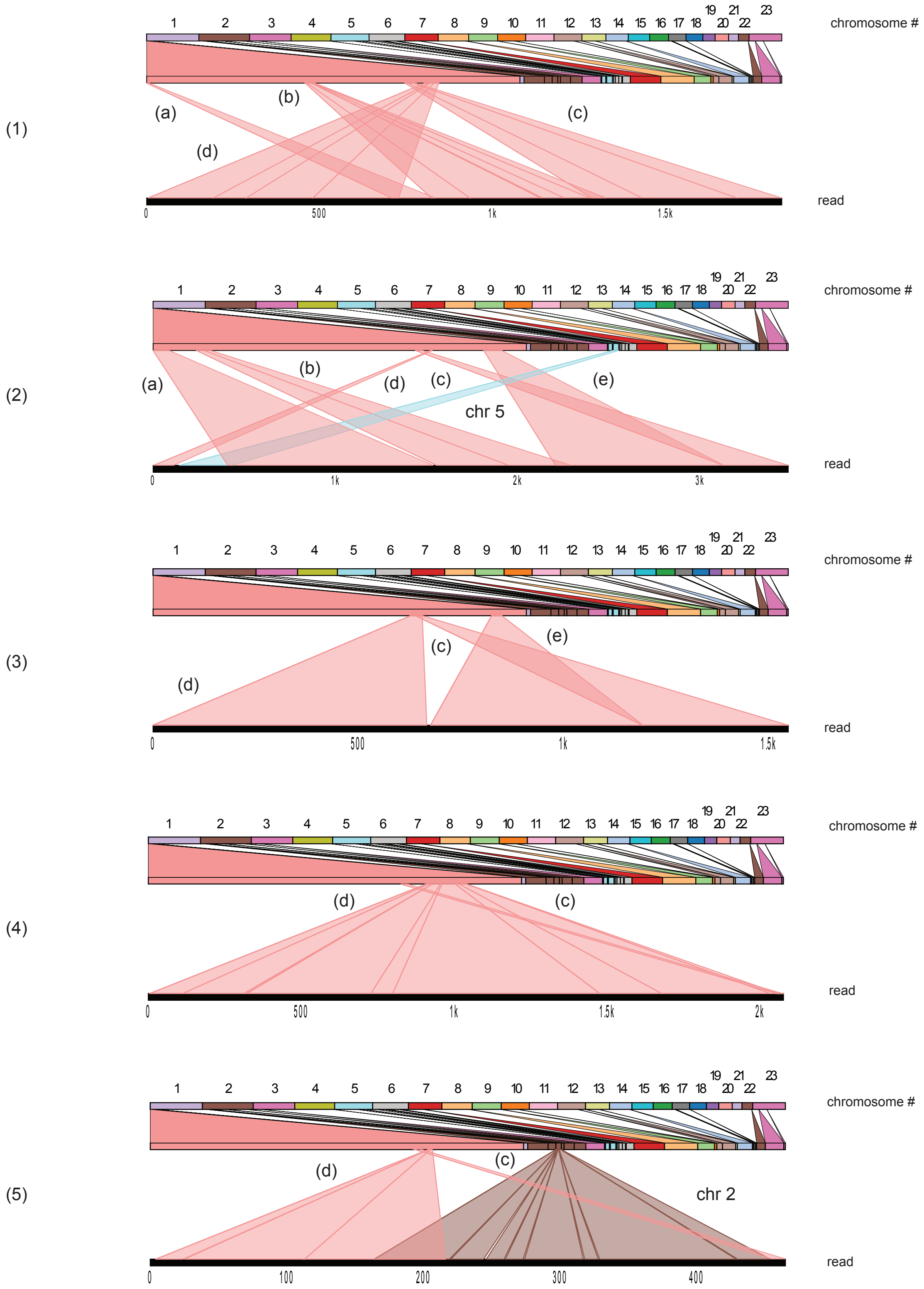

Supplementary Figure 7C

### $(CAG)_{102}$ clone 13 plus APH

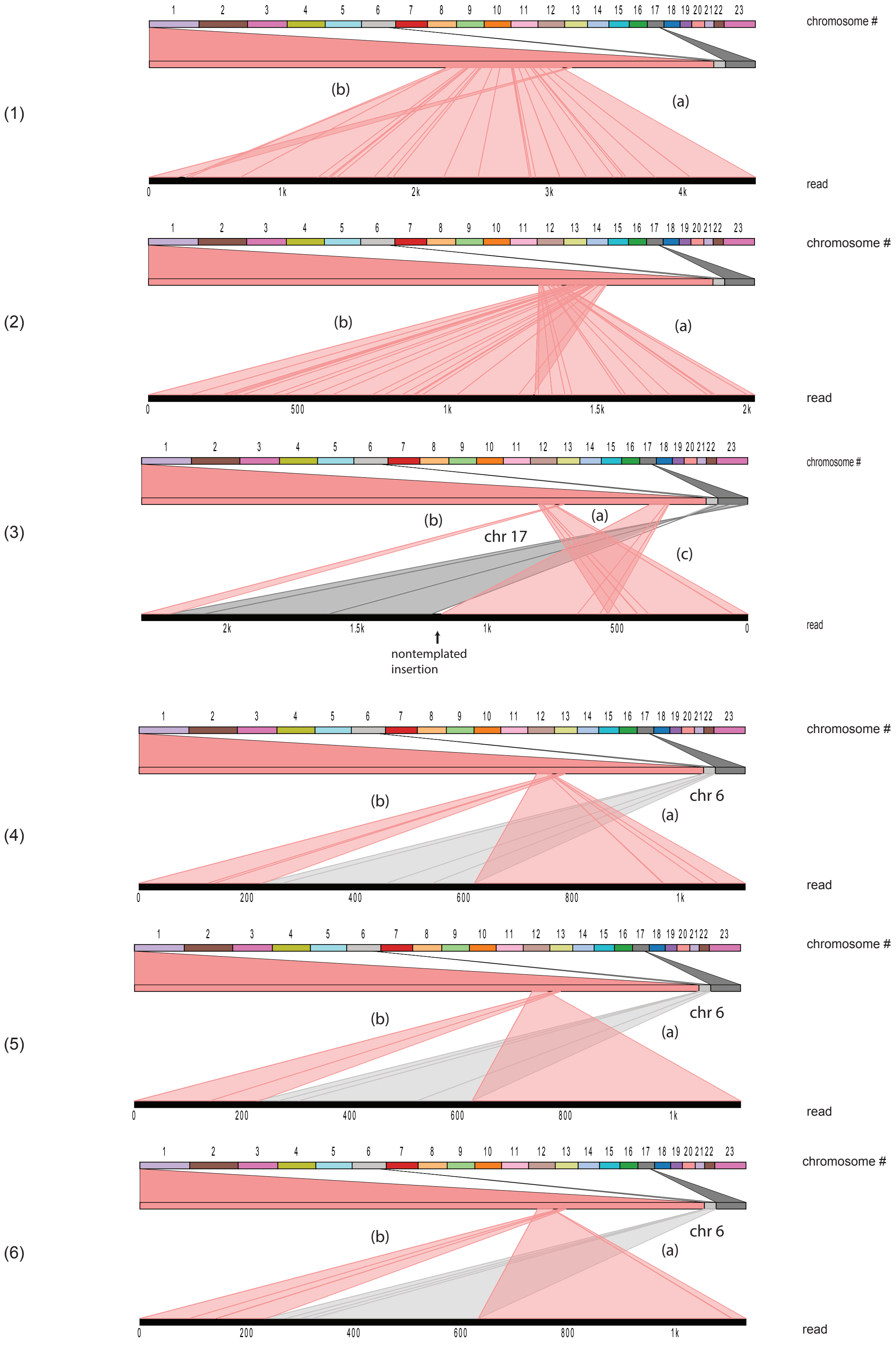

Supplementary Figure 7D

(CAG)$_{102}$ clone 13 plus APH

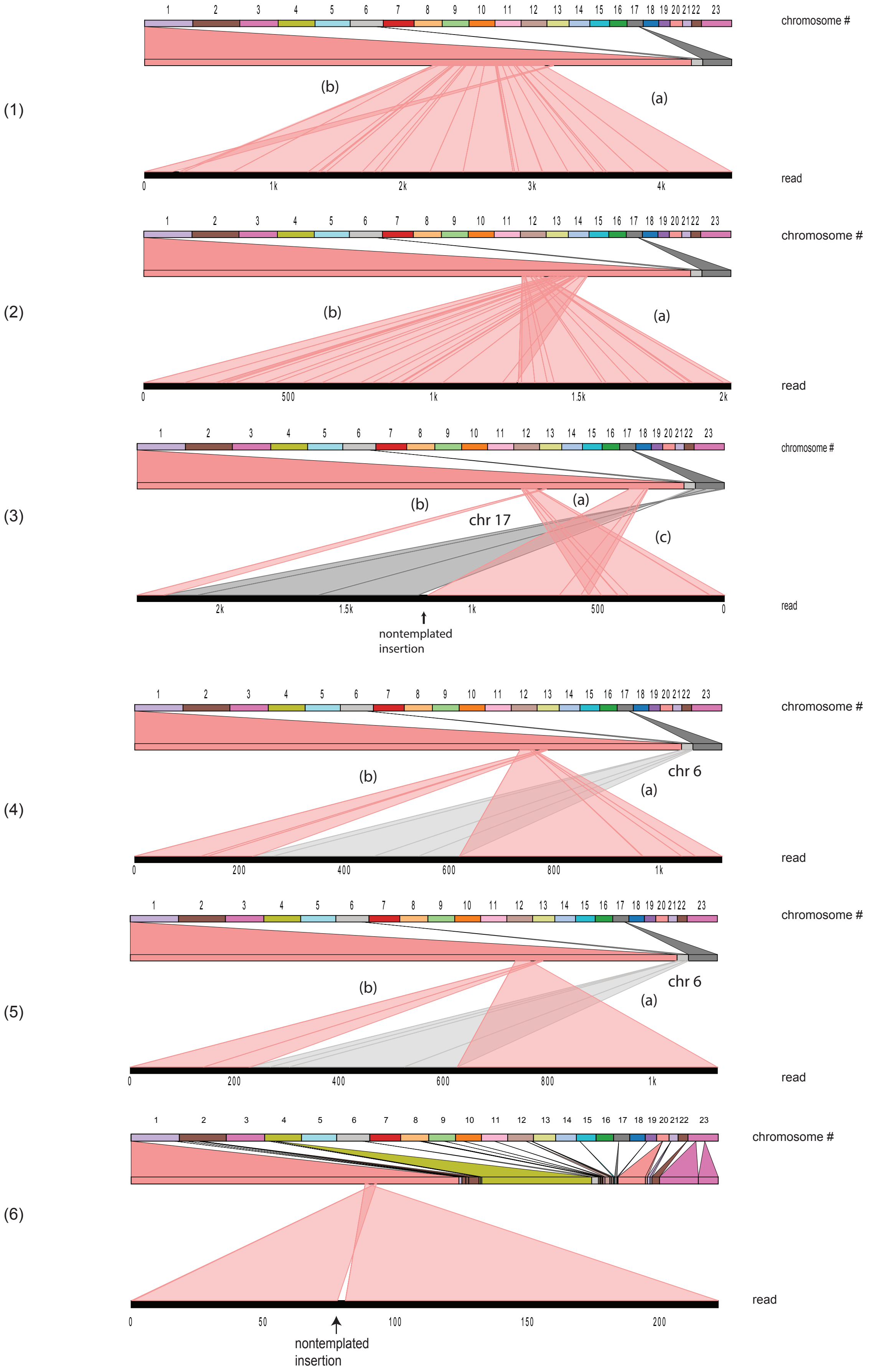

**Supplementary Figure 7: Template switching patterns of inhibitor treated (CAG)102 cells. (A) Query viewport view of reads from (CAG)102 c.10 cells treated with HU. (B) Reads from (CAG)102 c.10 cells treated with APH. (C) Reads from (CAG)102 c.13 cells treated with HU. (D) Reads from (CAG)102 c.13 cells treated with HU. Chromosome numbers are shown at the top; the lower heavy black line in each panel is the complete read. Heavier red lines indicate indels.**
