## Supplementary Figure 8 for "Microsatellite break-induced replication generates highly mutagenized extrachromosomal circular DNAs"

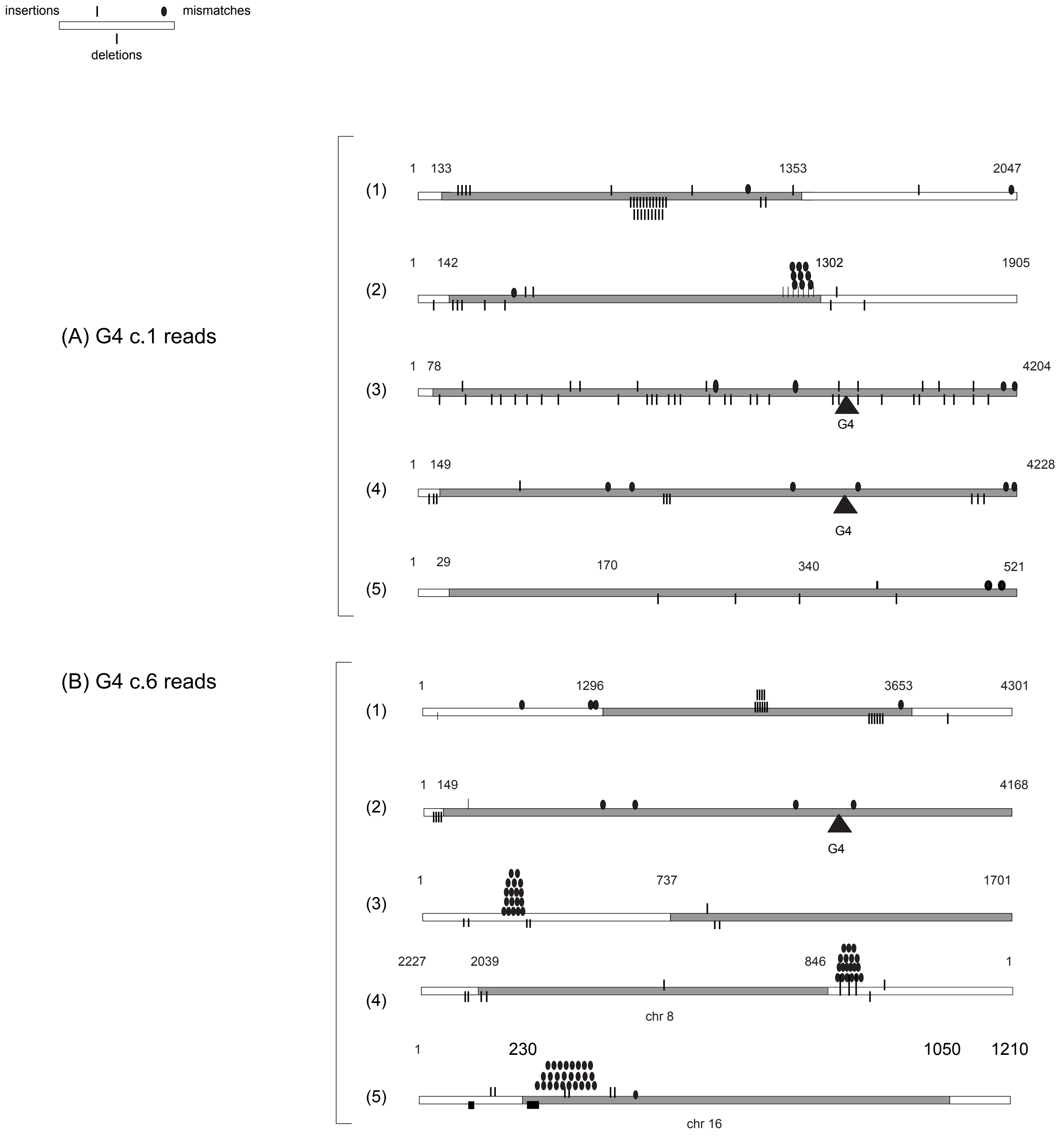

Supplementary Figure 8
*(continued)*

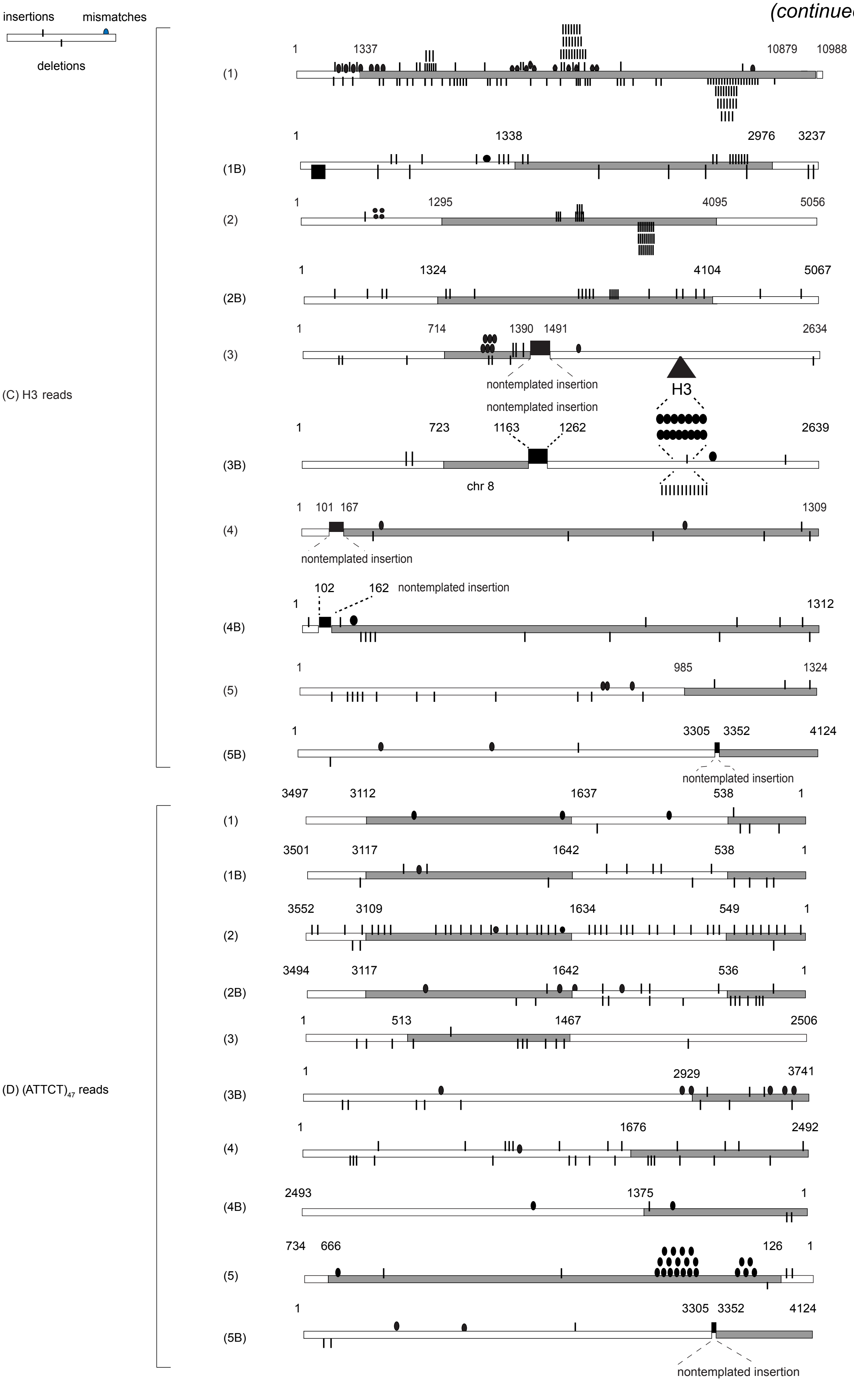

(E)

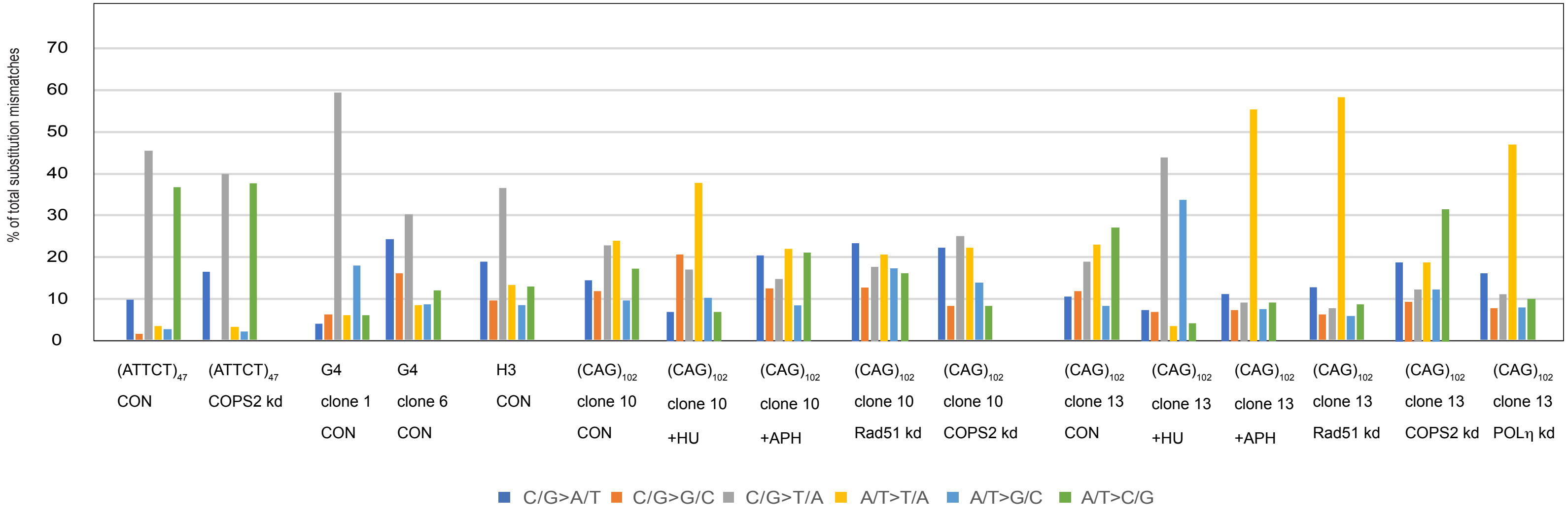
% of total substitution mismatches
70
60
50
40
30
20
10
0
(ATTCT)47 CON
(ATTCT)47 COPS2 kd
G4 clone 1 CON
G4 clone 6 CON
H3 CON
(CAG)102 clone 10 CON
(CAG)102 clone 10 +HU
(CAG)102 clone 10 +APH
(CAG)102 clone 10 Rad51 kd
(CAG)102 clone 10 COPS2 kd
(CAG)102 clone 13 CON
(CAG)102 clone 13 +HU
(CAG)102 clone 13 +APH
(CAG)102 clone 13 Rad51 kd
(CAG)102 clone 13 COPS2 kd
(CAG)102 clone 13 POLη kd
C/G>A/T
C/G>G/C
C/G>T/A
A/T>T/A
A/T>G/C
A/T>C/G

(F)

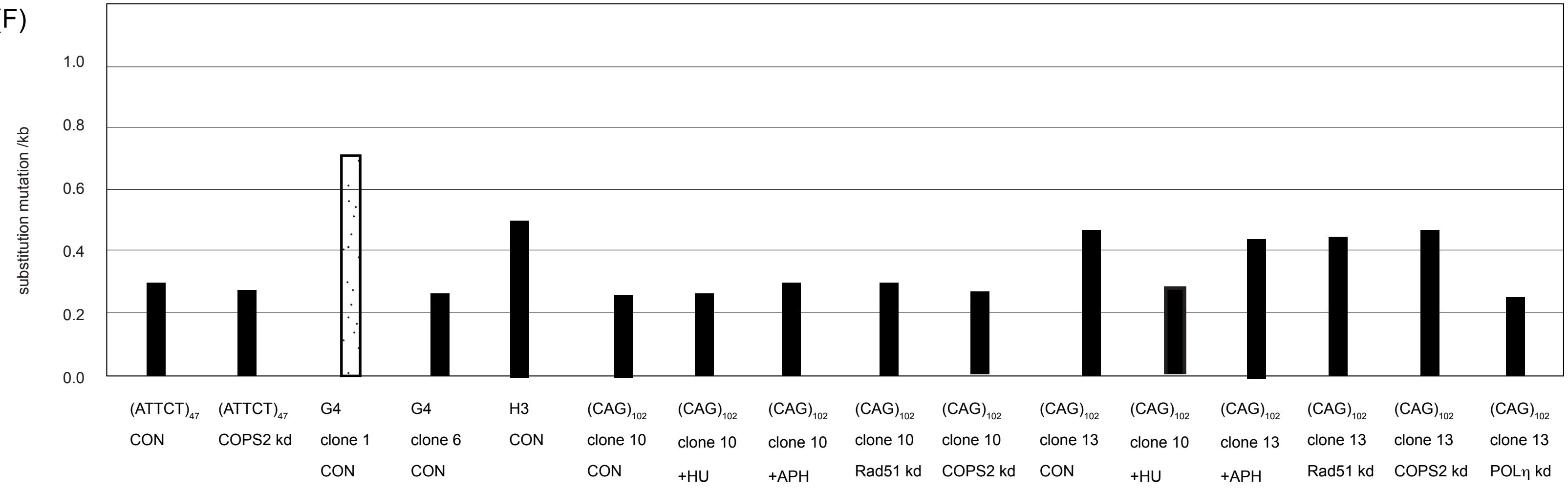
substitution mutation /kb
1.0
0.8
0.6
0.4
0.2
0.0
(ATTCT)47 CON
(ATTCT)47 COPS2 kd
G4 clone 1 CON
G4 clone 6 CON
H3 CON
(CAG)102 clone 10 CON
(CAG)102 clone 10 +HU
(CAG)102 clone 10 +APH
(CAG)102 clone 10 Rad51 kd
(CAG)102 clone 10 COPS2 kd
(CAG)102 clone 13 CON
(CAG)102 clone 10 +HU
(CAG)102 clone 13 +APH
(CAG)102 clone 13 Rad51 kd
(CAG)102 clone 13 COPS2 kd
(CAG)102 clone 13 POLη kd

**Supplementary Figure 8. Analysis of mutations.** The approximate positions of mutations were mapped in BLAST for (A) G4 c.1 read lines (1)-(5) (Figure 4(C)); (B) G4 c.6 read lines (1)-(5) (Figure 4(D)); (C) H3 cell read lines (1)-(5) (Figure 4(E)) and neighboring reads (1B-5B); (D) (ATTCT)47 cell read lines (1)-(5) (Figure 4(F)) and neighboring reads (1B-5B). (E) The mutational signatures of the eccDNAs from (CAG)102, G4, H3, and (ATTCT)47 cell lines, as well as HU treated, and APH treated (CAG)102 cells, are tabulated. (F) Base substitution frequencies of the samples shown in (A)-(E). Note the signature and mutation rate of G4 clone 1 cells (stippled), dominated by template switches with distinct end points, to a short region of chr14 (nt 91,293,050 - 91,293,650) containing eight G4 consensus matches within 600 bp.
