## Supplementary Table 1, 2 for "Microsatellite break-induced replication generates highly mutagenized extrachromosomal circular DNAs"

Supplementary Table 1: si/shRNAs

| | |
|---|---|
| siPOLη (SMARTpool) | AAACUGGCCUGAGGA |
| | CUAAGAAGUUAUGUCCAGAUCUU |
| | GCACUUACAUUGAAGGGUU |
| | GCAAUUAGCCCAGGAACUA |
| siCOPS2 | CAAGACGAACCACUUGCUUAA |
| shRad51 | CGCCCUUUACAGAACAGACUACUCGAGUAGUCUGUUCUGUAAAGGGCG |

Supplementary Table 2: iPCR primers

| Sample name | Forward primer | Reverse primer |
|---|---|---|
| $(CAG)_{102}$ c.10 | ATGTCCCGTCTGTTGTGTGACTCT | CGCTGCCGTCCTCGATGTTG |
| $(CAG)_{102}$ c.13 | AAGCTTGCCTTGAGTGCTTC | AATTGTCCATGCCGAGAGTGATC |
| $(ATTCT)_{47}$ | GGTATCTGTTTTCTATTTGTCTTCGGGAG | AGAATAGAATTTTTGAGATGAAGTCTCT |
| G4 c.1 | AACACCTAAAGCTTGCCTTGAGTGCTTC | AACACCTAGTCCATGCCGAGAGTGATC |
| G4 c.6 | ATATAGGAAAGCTTGCCTTGAGTGCTTC | ATATAGGAGTCCATGCCGAGAGTGATC |
| H3 | ACGTAGCTAAGCTTGCCTTGAGTGCTTC | ACGTAGCTGTCCATGCCGAGAGTGATC |
